# Chromosome structural rearrangements in invasive haplodiploid ambrosia beetles revealed by the genomes of *Euwallacea fornicatus* and *Euwallacea similis* (Coleoptera, Curculionidae, Scolytinae)

**DOI:** 10.1101/2024.06.25.600716

**Authors:** James R. M. Bickerstaff, Tom Walsh, Leon Court, Gunjan Pandey, Kylie Ireland, David Cousins, Valerie Caron, Thomas Wallenius, Adam Slipinski, Rahul Rane, Hermes E. Escalona

## Abstract

Bark and ambrosia beetles are among the most ecologically and economically damaging introduced plant pests worldwide, with life history traits including polyphagy, haplodiploidy, inbreeding polygyny and symbiosis with fungi contributing to their dispersal and impact. Species vary in host tree ecologies, with many attacking stressed or recently dead trees, such as the globally distributed *E. similis* (Ferrari). Other species, like the Polyphagous Shot Hole Borer (PSHB) *Euwallacea fornicatus* (Eichhoff), can attack over 680 host plants and is causing considerable economic damage in several countries worldwide. Despite their notoriety, publicly accessible genomic resources for *Euwallacea* Hopkins species are scarce, hampering better understanding of their invasive capabilities as well as modern control measures, surveillance and management. Using a combination of long and short read sequencing platforms we assembled and annotated high quality (BUSCO > 98% complete) chromosome level genomes for these species. Comparative macro-synteny analysis showed an increased number of chromosomes in the haplodiploid inbreeding species of *Euwallacea* compared to diploid outbred species, due to fission events. This suggests that life history traits can impact chromosome structure. Further, the genome of *E. fornicatus* had a higher relative proportion of repetitive elements, up to 17% more, than *E. similis*. Additionally, metagenomic assembly pipelines identified microbiota associated with both species including *Fusarium* fungal symbionts and a novel *Wolbachia* strain. These novel genomes of haplodiploid inbreeding species will contribute to the understanding of how life history traits are related to their evolution and will contribute to the management of these invasive pests.

**Significance:** Scolytinae are significant forestry pests around the world and commonly translocated due to human trade of wood and plant products. Life history traits including inbreeding and haplodiploidy are attributed to their successful establishment in novel environments. *Euwallacea fornicatus* is widely distributed and attacks a wide variety of live host trees. This study reports the genome of this species and for, *E. similis*, which colonises dead host trees. The genome assemblies presented herein are highly complete and scaffolded to pseudo-chromosomal level. Comparative analyses of these genomes and of other Scolytinae highlight significant chromosomal rearrangements between haplodiploid inbreeding *Euwallacea* and diploid outbreeding scolytinae species. Higher relative proportions of transposable elements were identified *E. fornicatus*, which may promote the species’ ability to attack live host trees. These genomes are the first for haplodiploid beetles and will contribute to the understanding of evolution of life history traits and the management of invasive insects.

## Introduction

Bark and ambrosia beetles (Curculionidae, Scolytinae) are some of the most ecologically damaging and economically important insect plant pests, contributing to tree mortality through infestation and the spread of pathogenic fungi (Hulcr & Stelinski 2017; Biedermann et al. 2019). Most bark beetles are phloeophagous and obtain nutrition from the phloem tissue while some lineages, termed ambrosia beetles, are xylomycetophagous and consume farmed symbiotic ambrosia fungi (Jordal 2014; Hulcr & Stelinski 2017). Many species are commonly transported to novel areas through human mediated movement of wood and plant products and over one hundred species have established outside of their native range (Lantschner et al. 2020). Life history traits including polyphagy, haplodiploidy, inbreeding polygyny and symbiosis with ambrosia fungi has been noted to contribute to the establishment of introduced species (Jordal et al. 2001; Lantschner et al. 2020; Vilardo et al. 2022), all of which characterise members of the tribe Xylebroini. Consequently, many such species have a global native or introduced distribution and are high priority pests in many regions of the world.

The genus *Euwallacea* Hopkins include several well studied polyphagous tree pests (O’Donnell et al. 2016). The genus is composed of over 80 species which are mostly circumtropical in their distribution (Storer et al. 2015). Species vary in their ecological relationships with host trees, with many species colonising stressed, cut and recently dead hosts. One such example is *E. similis* (Ferrari) (Figure 1b), a species native to South East Asia and possibly to Australia (Beaver et al. 2014), that has been introduced to the Americas and tropical African countries (Gomez et al. 2018) with 145 known host plants (Ruzzier et al. 2023). *Euwallacea fornicatus* (Eichhoff) (Figure 1a), also known as the Polyphagous Shot Hole Borer (PSHB) is considered one the most damaging Xyleborini species. It can attack live hosts and is known to attack over 680 tree species (Gomez et al. 2019; Mendel et al. 2021; Ruzzier et al. 2023) and is able to successfully reproduce in 168 species (Mendel et al. 2021; Ruzzier et al. 2023). *Euwallacea fornicatus* is widely distributed and has been introduced to California (Eskalen et al. 2012), Hawaii (Rugman-Jones et al. 2020), South Africa (Paap et al. 2018), Israel (Mendel et al. 2012), Europe (Schuler et al. 2023) and, recently, in the city of Perth, Western Australia (WA) where it is currently subject to an eradication campaign. This ambrosia beetle species is associated with several pathogenic *Fusarium* fungi, including *F. euwallaceae* Freeman *et al*. (2013) and *F. sp.* AF-18, and these fungi can induce wilt and dieback in host trees leading to tree mortality (Freeman et al. 2013). Economic modelling has estimated the damages caused by this species to $18.45 billion over a ten-year period in South Africa (De Wit et al. 2022) while in Perth WA the cost of eradicating this species is estimated to be AUD$45 million and management of the species could reach AUD$6.8 million per annum in 30 years (Cook & Broughton 2023).

**Fig 1.**
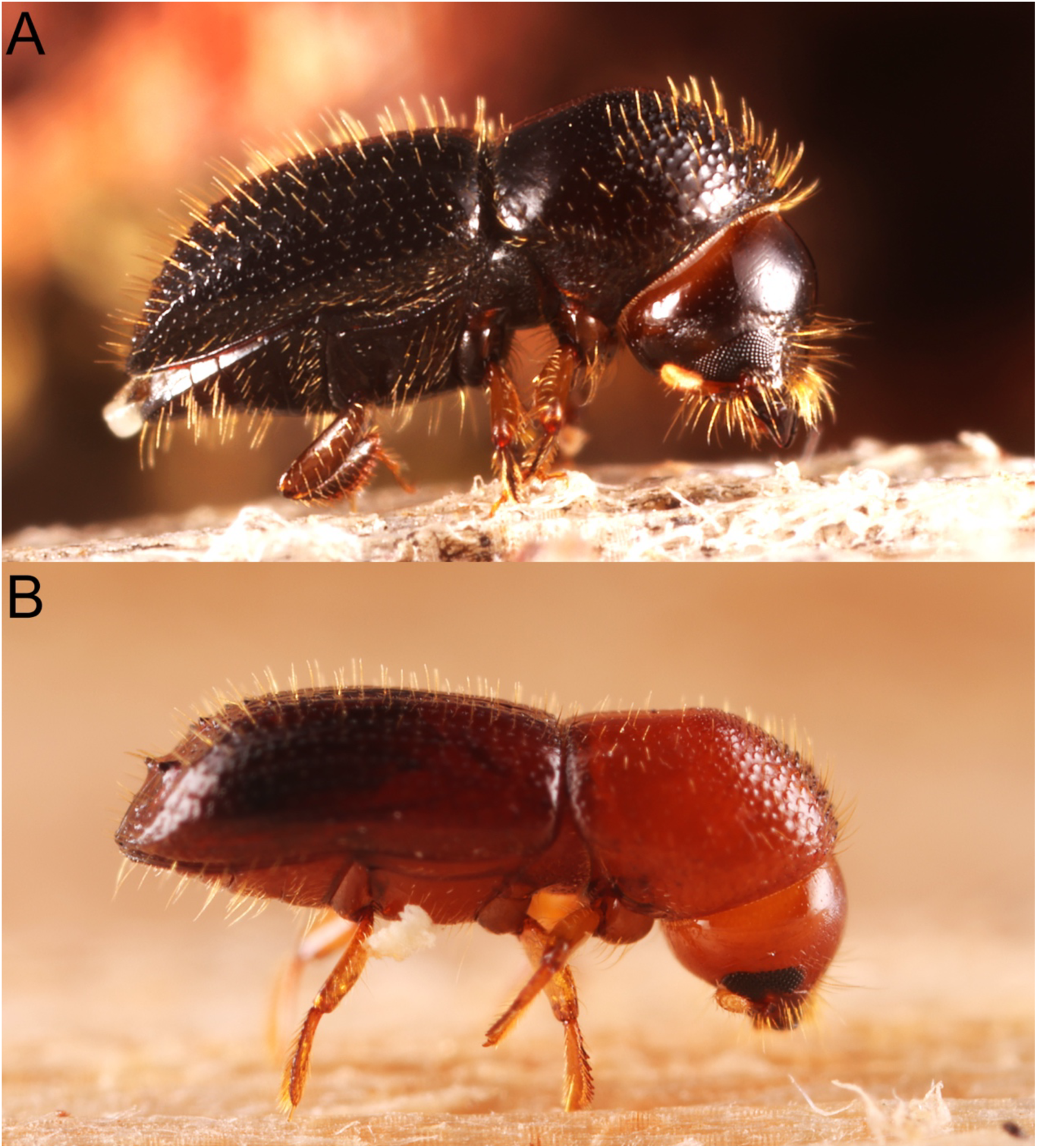
The ambrosia beetles A, *Euwallacea fornicatus* (PSHB) and B, *Euwallacea similis* whose genomes were sequenced herein. Photo taken by Hermes E. Escalona.

The species *E. fornicatus* has been associated with the species *E. perbrevis* (Schedl), the Tea Shot Hole Borer (TSHBa), *E. fornicatior* (Eggers), also the Tea Shot Hole Borer (TSHBb) and *E. kuroshiro* (Gomez and Hulcr), the Kuroshio Shot Hole Borer (KSHB) representing a species complex with similar morphology (Gomez et al. 2018; Smith et al. 2019) . Originally, these four species were all considered to be *E. fornicatus*, however, recent molecular phylogenetic and taxonomic works have stabilised their species status (Stouthamer et al. 2017; Gomez et al. 2018; Smith et al. 2019). Few genomic resources exist for *E. fornicatus* and other Xyleborini, with much of the data available restricted to mitochondrial *cytochrome oxidase I* (*COI*) and a handful of genes for phylogenetic analyses (Stouthamer et al. 2017; Gomez et al. 2018; Bierman et al. 2022). Mitochondrial *COI* phylogenies have identified 21 haplotypes of *E. fornicatus* with much of the diversity found in Vietnam and Taiwan (Stouthamer et al. 2017). One haplotype, H33, has been identified in California, South Africa and Israel and is thought to have originated from Vietnam (Stouthamer et al. 2017; Bierman et al. 2022) however, other haplotypes, H35 and H38, have also been found in California and South Africa, respectively, indicating that the species could have been introduced multiple times to both countries (Stouthamer et al. 2017; Bierman et al. 2022). To date, the only sequence data available for *E. similis* is restricted to *COI* barcodes in its native South East Asian distribution (Lynn et al. 2021).

Genomic data are scarce for these widespread pests and further molecular resources are vital to understand the invasion patterns of *Euwallacea* species, their invasive capabilities, and exploring novel ways of controlling outbreaks and mitigating future harm. Access to high quality genomes has led to advancements in the management of invasive species, through facilitating accurate and time-efficient diagnostics and surveillance (Frey et al. 2022; McGaughran et al. 2024), understanding the origin of invasive species and invasion pathways (Rane et al. 2023) and identifying genetic and genomic variation related to invasion potential (Sillero et al. 2020; McGaughran et al. 2024). In this study we present chromosome-scale, annotated genomes of *E. fornicatus* and *E. similis* using newly developed long read sequencing technologies and performed comparative analysis of macrosynteny, transposable elements, metagenomics and nuclear and mitochondrial phylogenetics. These genomes will be valuable assets for tracking the invasion pathway, developing novel biosecurity management plans and will provide essential insight into comparative of invasive species genomics.

## Results

### Genome assemblies and evaluation

In total 98.23Gb of high-quality Pacific Biosciences High Fidelity (PacBio HiFi) long sequence reads with a mean N50 of 11.6Kb were obtained to assemble four genomes of two *Euwallacea* species (Table S1). Two libraries of both species were sequenced and are herein referred to as EFF26 and EFF42 (*E. fornicatus*) and ESF13 and ESF15 (*E. similis*). The estimated genome size for *E. fornicatus* was between 192.7 Mb and 208.9 Mb, while estimates of 170.7 Mb and 181.8 Mb were made for *E. similis* and all are predicted to be highly homozygous (het: 0.012% – 0.112%), and all libraries consistently predicted similar results (Figure S1). All genomes were assembled with HiCanu (Nurk et al. 2020) as these assemblies closer to the estimated genome size compared with hifiasm (Cheng et al. 2021), and assembly quality metrics were similar between assemblers. Genomes were assembled with, post sequence read filtering and subsampling, 6.5Gb for EFF26 (29.58x, 0.35x subsample), 7Gb for EFF42 (24.07x, 0.35x subsample), 5.63Gb for ESF13 (30.95x, not subsampled) and 8.09Gb for ESF15 (44.78x, 0.15x subsample) of PacBio HiFi reads and 3.02Gb (*E. fornicatus*) and 3.3Gb (*E. similis*) of Arima Hi-C paired-end reads (Table S1).

Scaffolding was performed with both YahS (Zhou et al. 2023) and SALSA2 (Ghurye et al. 2019). While both scaffolders identified a similar number of pseudochromosome scaffolds for the *E. fornicatus* assemblies, SALSA2 failed to break incorrectly joined contigs identified through Hi-C read mapping for the *E. simils* scaffolds (Figures S2 – S5). As such, we present the pseudo-chromosomal scaffolded assemblies produced through YahS followed by decontamination with the BlobToolKit (Challis et al. 2020) and manual curation. In total, 39 pseudo-chromosomes were identified for the *E*.

*fornicatus* (EFF26, Figure S6; and EFF42, Figure S7) assemblies. Both EFF26 and EFF42 assemblies were larger than the estimated genome sizes, at 219.83 Mb and 239.6 Mb, respectively. For both *E. similis* assemblies 37 pseudo-chromosomes were identified (ESF13, Figure S8; and ESF15, Figure SS9). The ESF13 assembly was smaller than the estimated genome size, at 177.6 Mb, while the ESF15 genome was larger than estimated at 180.07 Mb. All assemblies were found to be highly complete and contained over 98% complete single copy BUSCO sequences (Table 1, Figure S10).

**Table 1.**
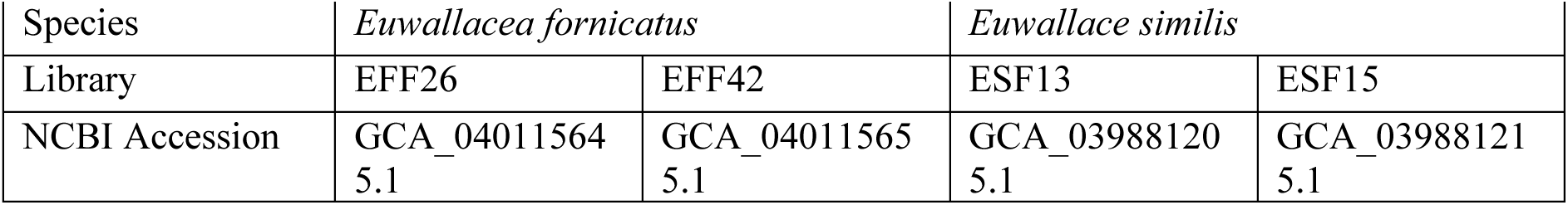
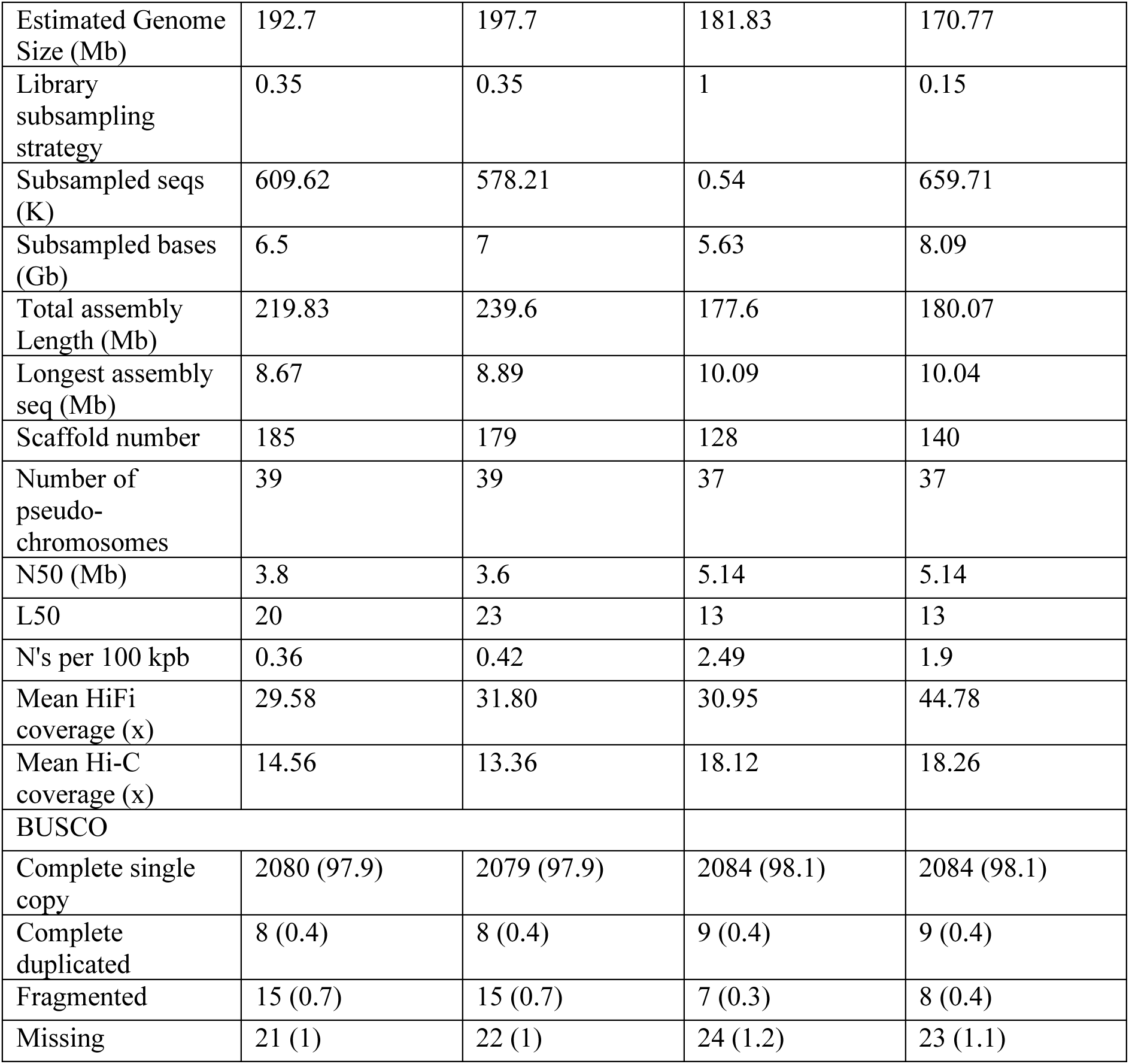
Genome summary statistics for the assemblies produced. Library subsampling strategy refers to the level of subsampling. Mean HiFi coverage was calculated by dividing the number of base pairs in the filtered subsampled HiFi libraries by the total assembly length. Mean Hi-C coverage was calculated by dividing the number of base pairs in the filtered Hi-C libraries by the total assembly length.

### Genome annotation

Genome annotation with the NCBI Eukaryotic Annotation Pipeline predicted 24,417 transcripts and 12,360 genes for the EFF26 *E. fornicatus* genome. Annotation for the ESF13 *E. similis* genome resulted in 19,838 transcripts and 11,978 genes predicted. Protein coding genes occupied 46 Mb of the *E. fornicatus* genome and 38 Mb of *E. similis,* 21% of both assembly lengths, and their density varied across chromosomes for both species (Figure 2). Annotations were largely complete with >97% complete single copy BUSCOs (Table 2). The number of genes annotated for both *Euwallacea* species were lower than those annotated for all other Scolytinae species, roughly 17 – 81% fewer (Navarro-Escalante et al. 2021; Powell et al. 2021; Liu et al. 2022; Wang et al. 2023).

**Fig 2.**
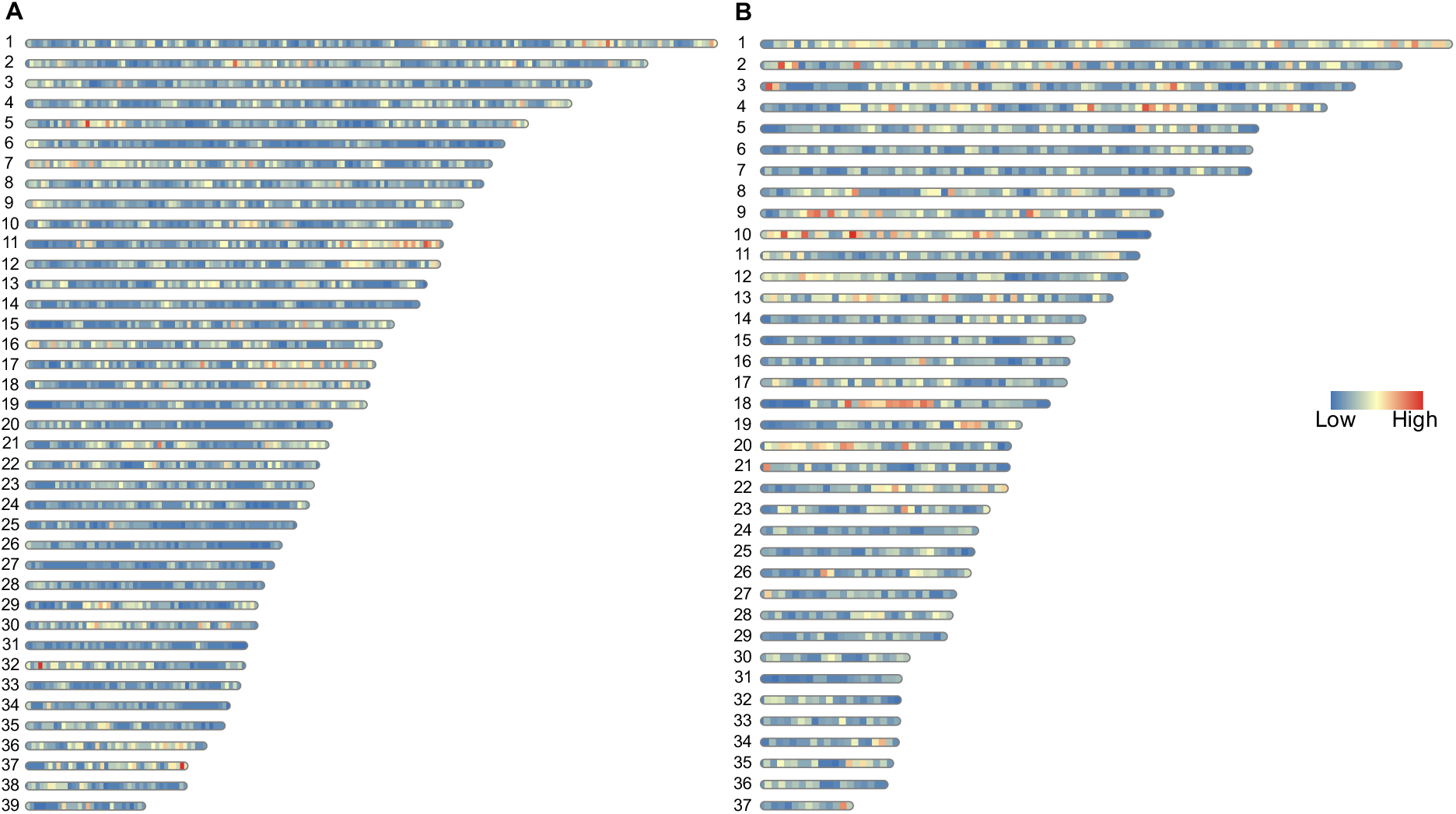
Distribution and density of annotated protein-coding genes along the pseudo-chromosome scaffolds for A, *Euwallacea fornicatus* and B, *Euwallacea similis*.

**Table 2.**
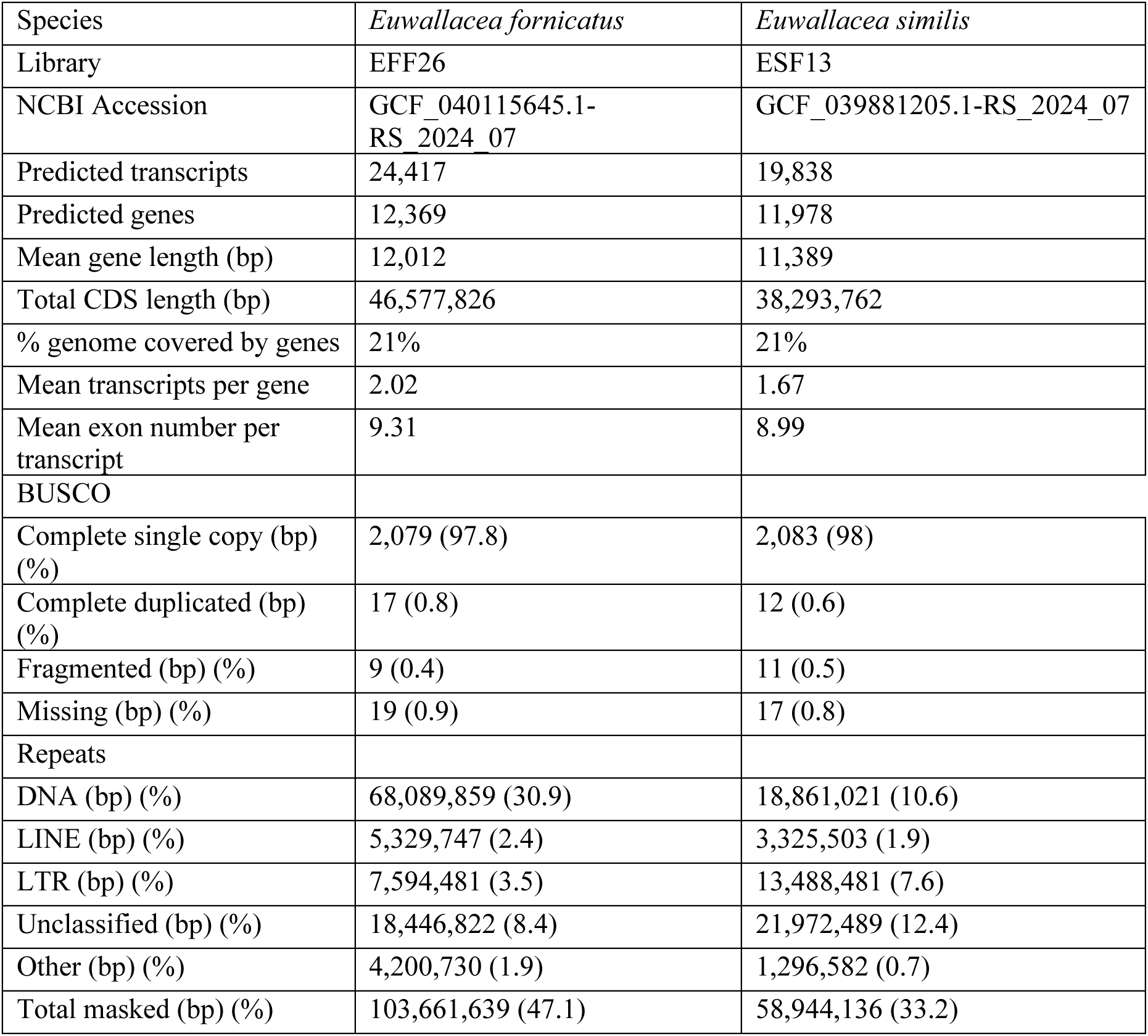
Protein and repeat annotation statistics for the assemblies produced. Proteins were identified and annotated by the NCBI Eukaryotic Annotation Pipeline and completeness assessed with BUSCO. Repeats were identified by EarlGreyTE.

| Species | *Euwallacea fornicatus* | *Euwallacea similis* |
| --- | --- | --- |
| Library | EFF26 | ESF13 |
| NCBI Accession | GCF_040115645.1-RS_2024_07 | GCF_039881205.1-RS_2024_07 |
| Predicted transcripts | 24,417 | 19,838 |
| Predicted genes | 12,369 | 11,978 |
| Mean gene length (bp) | 12,012 | 11,389 |
| Total CDS length (bp) | 46,577,826 | 38,293,762 |
| % genome covered by genes | 21% | 21% |
| Mean transcripts per gene | 2.02 | 1.67 |
| Mean exon number per transcript | 9.31 | 8.99 |
| BUSCO |  |  |
| Complete single copy (bp) (%) | 2,079 (97.8) | 2,083 (98) |
| Complete duplicated (bp) (%) | 17 (0.8) | 12 (0.6) |
| Fragmented (bp) (%) | 9 (0.4) | 11 (0.5) |
| Missing (bp) (%) | 19 (0.9) | 17 (0.8) |
| Repeats |  |  |
| DNA (bp) (%) | 68,089,859 (30.9) | 18,861,021 (10.6) |
| LINE (bp) (%) | 5,329,747 (2.4) | 3,325,503 (1.9) |
| LTR (bp) (%) | 7,594,481 (3.5) | 13,488,481 (7.6) |
| Unclassified (bp) (%) | 18,446,822 (8.4) | 21,972,489 (12.4) |
| Other (bp) (%) | 4,200,730 (1.9) | 1,296,582 (0.7) |
| Total masked (bp) (%) | 103,661,639 (47.1) | 58,944,136 (33.2) |

### Mitochondrial and nuclear phylogenetics

Phylogenetic analysis of mitochondrial haplotypes identified that the population of *E. fornicatus* in Perth WA belong to haplotype H38, with the *COI* barcode identical to populations native in China and Vietnam, and also to the invasive population in South Africa. The H38 haplotype clustered with Haplotypes 37, 39 – 42, and were found to be phylogenetically distinct from the more prevalent invasive H33 haplotype (Figure S11). Phylogenomic analysis using BUSCOs of published Scolytinae genomes and those of the reference *Euwallacea* yielded a topology similar to that reported by Pistone *et al*. (Pistone et al. 2018) and all nodes were strongly supported (Figure 3).

**Fig 3.**
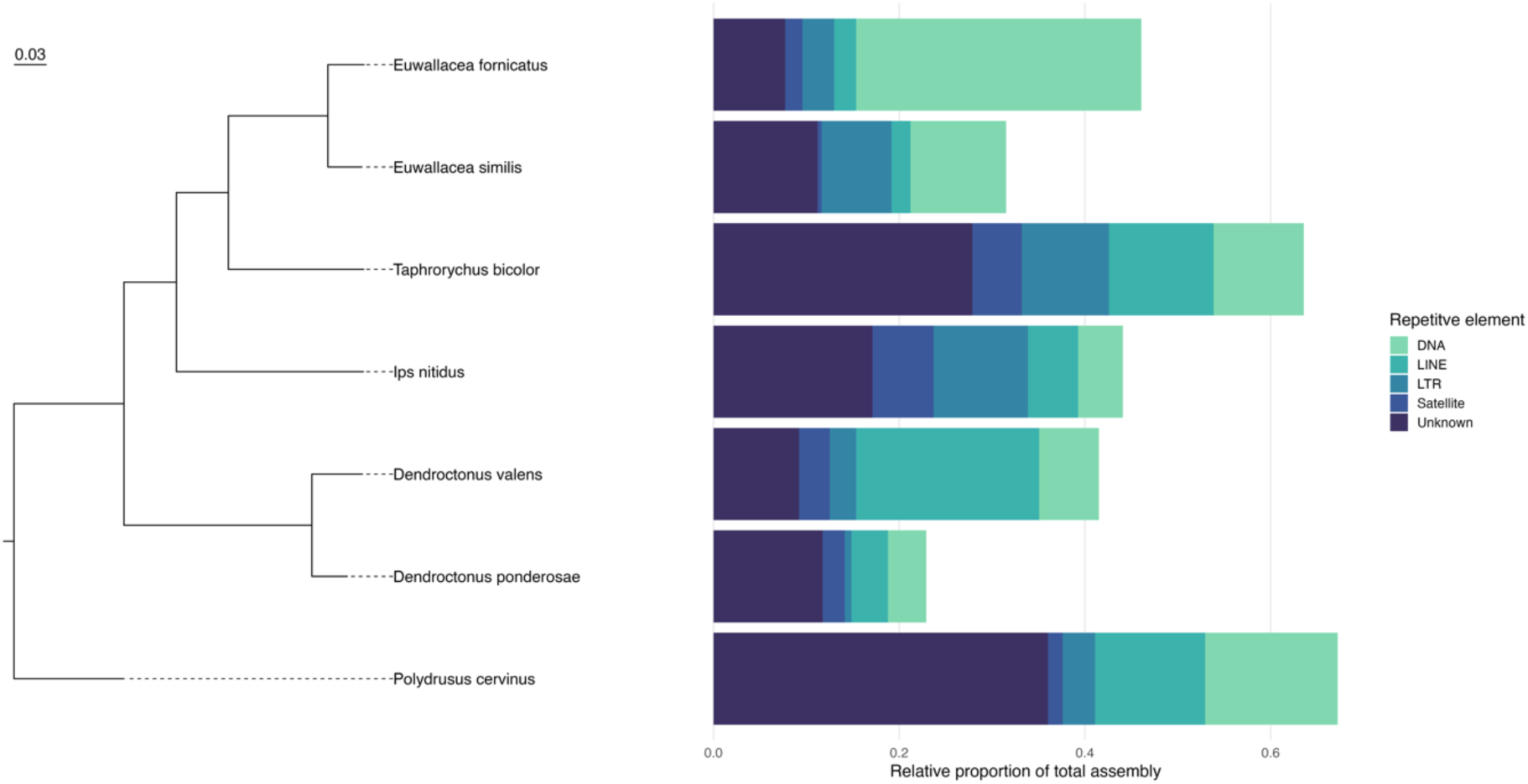
Maximum-likelihood phylogenetic tree pseudo-chromosome level assemblies for Scolytinae based using 2,108 single copy complete BUSCO sequences. All nodes were strongly supported with 100 bootstrap support. Scale bar indicated 0.03 substitutions per nucleotide position. To the right of the tips are the relative proportion of repetitive element composition as annotated in the species’ genome.

### Repeat element and chromosomal variation across Scolytinae

Comparative analyses of repeat and transposable elements found that their relative proportion in the genomes of Scolytinae varied across lineages. Roughly half (47 – 50%) of the *E. fornicatus* assemblies and over one third (33 - 34%) of the *E. similis* assemblies were masked by the EarlGreyTE pipeline (Table 2, Figure 2, Figure S12). DNA transposons and long terminal repeats (LTR) were the most abundant repeat elements identified, with unclassified elements only accounting for a small proportion of all annotations (Table 2). Few rolling circle elements were identified in all assemblies and SINEs were only found in the *E. similis* assemblies. Kimura 2 divergence landscapes of repetitive elements indicate a recent burst of DNA transposons and LTR elements, while LINE and rolling circle elements are experiencing ongoing proliferation in the *Euwallacea* assemblies (Figures S13 – 16). The relative proportion of annotated TEs varied throughout Scolytinae assemblies, with over 60% of the *T. bicolor* assembly occupied by TEs, and the assembly of *D. ponderosae* possessed the lowest relative proportion of annotated TEs (25%). The relative proportions of the types of TEs also varied significantly throughout Scolytinae as well, and TE annotations were largely composed of unclassified elements, DNA transposons and LINEs. All published Scolytinae assemblies possessed greater amounts of Satellite sequences than the *Euwallacea* assemblies (Figure 3). Lastly, the Entiminae outgroup *P. cervinus* had a larger proportion of the genome of annotated TEs compared to Scolytinae.

The chromosome number and synteny of haplodiploid inbreeding *Euwallacea* species greatly varied compared to diploid outbreed Scolytinae species. The assemblies of *E*. *fornicatus* and *E. similis* were more syntenic with one another than other Scolytinae (Figure S17). Largely, synteny was retained throughout the Scolytinae assemblies with several translocations of genomic material. However, extensive translocations, rearrangements, and fissions of syntenic blocks were identified between the assemblies of *Euwallacea* and other Scolytinae. Further, the X sex-chromosome was identified to be synentic throughout all assemblies, however, blocks corresponding to these scaffolds were broken apart in the *Euwallacea* assemblies (Figure 4).

**Fig 4.**
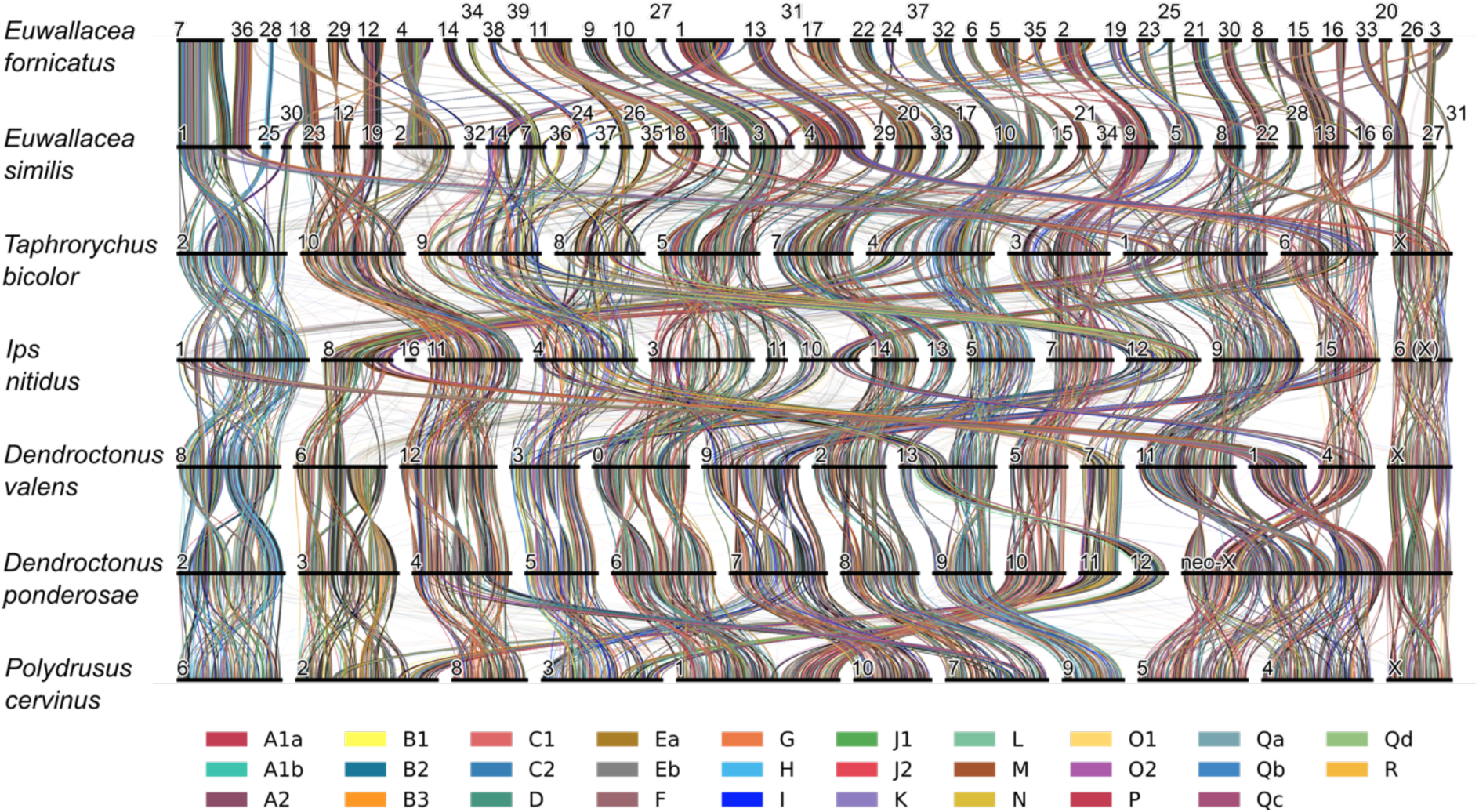
Ribbon plot showing the synteny and structural and chromosomal rearrangement in Scolytinae chromosomal assemblies. Syntenic blocks of single copy complete BUSCO sequences link pseudo-chromosome scaffolds between assemblies. Black bars represent individual pseudo-chromosome scaffolds and are labelled according to the name provided in the assembly. Scaffolds are ordered following the *Dendroctonus ponderosae* assembly order and the lengths of the bars representing scaffolds indicate total number of BUSCOs and not scaffold length. Pseudo-chromosome scaffold labels follow how they were published either on NCBI or their respective publications, the X-chromosome label in parentheses is inferred based on syntenic linkages. Coloured lines connect orthologous genes across scaffolds and the colour code follows the Bilaterian-Cnidarian-Sponge Linkage Groups (BCnS LGs) as in (Schultz et al. 2023).

### Metagenomics

The microbial diversity and composition varied between the two *Euwallacea* species. The two *E. similis* libraries produced more Metagenome-assembled genomes (MAG) contigs (730 and 1,200) than the two *E. fornicatus* libraries (103 and 109). Across all libraries, many more Bacteria were identified than other microbial Kingdoms, with Actinomycetota and Pseudomonadota highly prevalent (Figure S18). Several complete circularised Bacteria assemblies were also identified across all libraries (Table S2), including *Wolbachia* for both *E. similis* libraries. Multi-locus Sequence Typing (MLST) profiling identified this *Wolbachia* strain closely matching *w*Ei, which infects *E. interjectus* (Kawasaki et al. 2016), however there were SNPs in the *hcpA* and *gatB* loci. Consequently, this represents a novel strain, herein referred to as *w*Es. Several fungal taxa were identified for both species (Figure S18), and mitogenomic contigs of *Fusarium* belonging to the *Euwallacea Fusarium* Clade (EFC) were identified by BLAST for all libraries except for ESF13. The only fungi found in *E. fornicatus* MAGs were *Fusarium*, while a richer diversity of fungi were found in association with *E. similis*, with 13 genera classified including *Leptographium, Ophiostoma, Ceratocystis* and *Sporothrix* (Table S2). Several viral contigs were also identified in both species, all belonging to Caudoviricetes.

## Discussion

### Genome synteny across Scolytinae

We sequenced, assembled and annotated the genomes of *Euwallacea fornicatus* and *E. similis*. These genomes are highly contiguous, complete and accurate as we sequenced the libraries of two female specimens per species and employed multiple assembly and scaffolding pipelines to validate their quality. The *E. fornicatus* assemblies presented herein were accepted and released on the NCBI GenBank on the same day as two other contig-level assemblies of the same species from South Africa (GCA_040114445 & GCA_040114495), from a female and male, respectively. The assembly sizes of these contig-level genome assemblies are similar to the chromosome-scaffold assemblies presented here, further validating the quality of the assemblies presented herein. These genomes represent the first chromosome scaffold genomes for these pests and for a haplodiploid inbreeding beetle lineage. They are among the smallest Scolytinae genomes published and contain the highest number of pseudo-chromosomal scaffolds. Additionally, about twelve thousand protein coding genes were predicted for these *Euwallacea* assemblies, representing 17 – 81% fewer proteins compared to previously published Scolytinae genomes (Navarro-Escalante et al. 2021; Powell et al. 2021; Liu et al. 2022; Wang et al. 2023).

The number of pseudo-chromosomal scaffolds identified for both *Euwallacea* species (n = 37 – 39) were higher than the typical number of chromosomes in Scolytinae (n = 7 – 16) (Virkki & Smith 1978). Chromosome number variation has previously been identified across and within genera of scolytines. For example, the karyotype of all 19 *Dendroctonus* species have been characterised, with chromosome numbers ranging from 2n = 12 – 30, and chromosomal fusion resulting in chromosome number reduction (Lanier 1981; Six & Bracewell 2015). Similarly, chromosome number variation has also been identified across 32 species of *Ips*, with numbers ranging from 2n = 16 – 32 (Virkki & Smith 1978). In Curculionidae, a larger variation in chromosome numbers have been characterised in those that are parthenogenetic due to polyploidy (Virkki & Smith 1978; Blackmon & Demuth 2015). For example, parthenogenetic pseudogamy resulting in triploid females have been characterised in *Ips acuminatus* Gyllenhal and *I. tridens* Mannerheim (Lanier & Kirkendall 1986). Similar observations have been made for several species of obligate inbreeding haplodiploid Xyleborini (Takenouchi & Takagi 1967; Virkki & Smith 1978). Species of the genus *Xylosandrus* Reitter have fewer chromosomes than other Scolytinae, with *X. germanus* (Blandford) 2n = 8 and *X. compactus* (Eichhoff) 2n = 10 (Takenouchi & Takagi 1967). In contrast, the chromosomes of *Anisandrus dispar* (F.) are small in size and numerous, n = 40 (Virkki & Smith 1978), similar to the *Euwallacea* species genomes.

We identified extensive chromosomal rearrangements between haplodiploid inbreeding and diploid outbreeding Scolytinae. The synteny of autosomal chromosomes and the ancestrally conserved X chromosome between diploid outbreeding species are largely conserved across genera, with syntenic blocks either translocated within or across scaffolds. Further, despite the difference in chromosomal numbers between *D. ponderosae* (2n = 12) and *D. valens* (2n = 14) (Six & Bracewell 2015), synteny across autosomal and X-chromosomes are largely conserved, identified herein and previously (Liu et al. 2022). Furthermore, extensive chromosomal rearrangement characterised the variation in genomic architecture between *Euwallacea* and diploid outbreeding species, with widespread translocation events shuffling syntenic blocks and fission events breaking up autosomal chromosomes. The conserved X chromosome was also found to be broken up through fission in the *Euwallacea* assemblies. This may relate to the apparent lack of sex chromosomes in other Xyleborini species as observed by Takenouchi & Takagi (Takenouchi & Takagi 1967) in male germ cells. Comparative analyses of Coleopteran karyotypes have identified that small effective population sizes and genetic drift greatly impact chromosome number variation across species (Blackmon et al. 2024). We speculate that the low genetic diversity and high homozygosity due to obligate inbreeding in *Euwallacea* may contribute to the increased chromosome number identified in these assemblies.

Comparative analyses indicate that the *E. fornicatus* genome has one of the highest proportions of repeat and transposable elements relative to total assembly among sequenced Scolytinae species, with roughly 50% of the assembly composed by TEs. While the relative proportion of these elements in the genome of *E. similis* only occupy about 30% of the total assembly, DNA transposons are among the most prevalent transposable elements in both *Euwallacea* species. DNA transposons have been identified to play a significant role in Eukaryotic genome and chromosomal evolution (Feschotte & Pritham 2007). Moreover, it has been identified in plants (Ren et al. 2018) that genomic rearrangement and recombination can result in net sequence loss at newly formed telomeric regions in the chromosome. Given that we have identified bursts in the proliferation of DNA transposons in the genomes of *Euwallacea*, these repetitive elements could be involved in the widespread structural variation, high number of chromosomes and smaller genome size and content in this genus.

Co-evolution with symbiotic ambrosia fungi could also contribute to chromosomal rearrangements in Xyleborini, as observed in *Euwallacea* assemblies. Genomic erosion is particularly prevalent in the genomes of symbionts that provide nutrition to their hosts, and genes of biosynthesis pathways producing amino acids obtained from the host are commonly lost (McCutcheon & Moran 2012; Sloan & Moran 2012). This has been identified to occur in host animals as well; in the deep-sea bone-eating worm, *Osedax frankpressi* Rouse, Goffredi & Vrjenhoek (Annelida: Polychaeta), endosymbionts provide key nutritional biosynethetic pathways which has subsequently resulted in host genome reduction and gene loss (Moggioli et al. 2023). Similarly, in socially parasitic ant species, gene loss of olfactory receptor repertoires corresponding with reduced social behavioural has evolved repeatedly (Schrader et al. 2021). Host genome rearrangements have also been reported in ants, with high levels of structural rearrangements, loss of synteny across assembled genomes, gene family contraction, expansion and a gain of novel genes identified in the *Atta* fungal farming ant lineage (Nygaard et al. 2011, 2016). Similar processes could have acted on the genomes of fungal farming Scolytinae in the coevolution with their symbionts, leading to the observed small genome size, genomic rearrangements and loss of synenty with other Scolytinae. It should be noted that the evolution of obligate inbreeding and haplodiploidy, in Dryocoetini, seems to precede the development of the ambrosia fungal symbiosis in Xyleborini (Jordal & Cognato 2012), and as such, further chromosome scale assemblies of non-fungus farming inbreeding haplodiploid species (Dryocoetini) are needed to assess the impact of the ambrosia symbiosis on chromosome structure.

### Genomic resources for biosecurity

The genomes assembled for *Euwallacea* will be essential resources for the management of these pests. A vital component in managing an invasive species is understanding where the species originates and how it arrived. Through mitochondrial haplotyping, we have identified that the population of *E. fornicatus* introduced to Perth, Western Australia corresponds to the H38 haplotype, which has also been introduced to California and South Africa. While further markers are needed for population genomic analyses in order to track the invasion pathway from the native to the introduced range (Rane et al. 2023), the assembled genomes will be vital in understanding population differentiation at the gene and genomic level. These genomes will also be a vital resource in the diagnostics and detection of these pests. While the mitochondrial genome for both species have already been published (Wang et al. 2020; Guo et al. 2023), the ability to identify nuclear markers or sequences will be important for the diagnostics and detection (such as through eDNA techniques) of the pests. Diagnostic protocols exist for the detection of *E. fornicatus* through identification of the symbiont *Fusarium euwallaceae* (De Jager & Roets 2022), however, the protocol operates on the detection of fungus in host plant woody tissues, which indicates that the species is already present and likely reproducing in a given locality. Using eDNA techniques and these reference genomes, these beetle pests could be detected early, increasing the chances of eradication or containment, and reducing the time and resources necessary to achieve this. Additionally, such methodologies could also demonstrate the absence of this pest in sampled areas allowing for cost effective determination of management and eradication outcomes.

Beyond tracking, detection and diagnostics, these genomes will also provide needed information for genome guided management of these pest. The endosymbiont gram-negative bacteria *Wolbachia* has been successfully deployed as a control agent for insect pests (Ross et al. 2022) and may prove useful in the control of *Euwallacea* pests. While strains have not yet been identified to be associated with *E. fornicatus*, further sequencing and MLST identification could identify naturally present strains, or strains from closely related species (such as *w*Es identified in this study) that could be artificially transferred (Xi et al. 2005). However, as noted in previous studies, the impacts of *Wolbachia* on haplodiploid inbreeding Scolytinae are not yet fully understood (Kawasaki et al. 2016; Bickerstaff et al. 2023). Gene drive through CRISPR gene editing may be an avenue to control populations of invasive *E. fornicatus* and *E similis.* As demonstrated in invasive wasps (Lester et al. 2020; Meiborg et al. 2023), the ability to control population growth with this technique is limited due to haplodiploidy, with only a single gene targeting female fertility identified as useful and infertile individuals were not as competitive as fertile counterparts. Outbreeding is rare in the inbreeding Scolytinae lineages and comes at a fitness cost (Peer & Taborsky 2005), and as such the penetration of gene drive through populations may be limited. Further insights are needed to first identify candidate genes in these genomes and then test the suitability of this management approach.

These genomes will also be of value in understanding mechanisms driving species invasion and pest impact from a genomic perspective. Preliminary comparative analyses of transposable element variation indicate that the relative proportion of genome content is similar to other beetle genomes (Petersen et al. 2019; Gilbert et al. 2021) however, a higher number of TEs are present in the *E. fornicatus* genome than in the *E. similis* genome. Transposable elements have been identified to be adaptive in species (Gilbert et al. 2021) and promote genetic diversity and rapid adaption in invasive species. The larger diversity of TEs present in the *E. fornicatus* genome may be associated with the species ability to infest a wide range of live host trees. Further comparative genomic analysis, such as gene family evolution (Powell et al. 2021), structural variation (Stapley et al. 2015; North et al. 2021), and pangenomics (Gerdol et al. 2020) of both species and their symbionts may yield greater insights into why these species are such successful invaders compared to other haplodiploid inbreeding ambrosia beetles, understanding variation in host tree ecology and why *E. fornicatus* is more destructive than other members *Euwallacea*.

## Conclusions

Pest species of *Euwallacea* are globally invasive and can cause significant ecological and economic impacts in the ranges where they are established. Our study presents the first chromosome-level genome assemblies of these pests and also for haplodiploid inbreeding beetles. These assemblies offer a deeper understanding of the genome structural variation related to species ecology and also serve as essential resources for biosecurity management and control strategies. The comparative analyses of repeat and transposable elements, as well as the macro-synteny of chromosome assemblies, shed light on the potential impacts of haplodiploidy and inbreeding on genome evolution in these beetles. Our findings underscore the importance of high-quality genome assemblies in advancing our knowledge of invasive species and contributing to their effective management and control.

## Material and methods

### Sample collection, DNA extraction and sequencing

Male and female specimens of *E. fornicatus* were collected by the Department of Primary Industries and Regional Development (DPIRD) WA from felled *Acer negundo* trees in Perth WA in October 2022. Male and females were flash frozen by storing them on dry ice and shipped to CSIRO Black Mountain Canberra Australian Capital Territory and stored at -80°C. *Euwallacea similis* larvae and adult females were collected from a fallen *Ficus* sp. branch in Eureka, northern New South Wales in April 2023. Specimens were transported from the field at 10°C and flash frozen by storing them at - 80°C. Vouchers of both species have been deposited in Australian National Insect Collection (ANIC) under the collection codes ANIC-25-086063 – ANIC-25-086073.

Female specimens were used for DNA extraction due to their larger size compared to haploid males. First, genomic DNA (for short read sequencing) of both species was extracted using a standard Qiagen DNEasy kit approach. We then used a single specimen from each species for high molecular weight DNA extraction with the Qiagen MagAttract kit following a modified protocol. Samples were incubated overnight at 56°C and lysates were agitated at 300 rpm. Following this, DNA was suspended in two elutions of 100 μl of Buffer EB. The concentration of all extracts were assessed using a DeNovix spectrophotometer and a Qubit Fluorometer with the Qubit dsDNA High Sensitivity Assay kit. Fragment size distribution and genomic DNA was assessed on an Aligent Fragment Analyzer system with the high sensitivity genomic DNA kit DNF-488.

For short read sequencing, sequencing libraries with 150 bp paired end reads were prepared and sequenced on the Illumina NovaSeq 6000 system at Genewiz (Suzhou, China). For long read sequencing, high molecular weight (HMW) DNA extracts were sequenced on the PacBio HiFi circular consensus Revio system at the AGRF in Brisbane, Queensland. For two HMW extracts of each species, gDNA was mechanically sheared, followed by SMRTbell Express 2.0 library construction. Short DNA fragments were removed using BluePippin. Barcoded libraries were then sequenced using one SMRT cell for each species on the PacBio Revio system; the libraries for *E. fornicatus* are herein referred to as EFF26 and EFF41, and the libraries for *E. similis* are herein referred to as ESF13 and ESF15.

### Genome size estimation, assembly and scaffolding

Genome size and heterozygosity estimates were performed with all sequence reads; kmers were counted with jellyfish (Marçais & Kingsford 2011) and genome size and heterozygosity were estimated with GenomeScope v2 (Ranallo-Benavidez et al. 2020). HiFi reads were then filtered for HiFi adapters with HiFiAdapterFilt (Sim et al. 2022) and subsampled with seqtk into randomly sampled subsets, ranging from 12x, 15x, 20x, 35x and 50x, of the estimated genome size, for all four libraries. Contigs were assembled with these subsets and the full long read dataset using HiCanu v2.1.1 (Nurk et al. 2020), with the genome size estimates of 190 Mb for both EFF26 and EFF42, and 170 Mb for both ESF13 and ESF15. Contigs were also assembled with hifiasm v0.19.6-r595 (Cheng et al. 2021) using the -l0 flag to disable purge duplication as these genomes are highly homozygous and presumably inbred. Misassembled contigs were assessed and corrected with CRAQ v1.0.9a (Li et al. 2023) and haplotigs and contig overlaps were removed with purge_dups v1.2.5 (Guan et al. 2020). The quality of the corrected contig assemblies were assessed with QUAST v5.2.0 (Mikheenko et al. 2018) and BUSCO v5.5.0 (Manni et al. 2021) using the endopterygota_odb10 dataset and the tribolium2012 dataset was specified for Augustus (Stanke et al. 2008) training.

Proximity ligation information was generated with Arima HiC 2.0 kits using tissue from a single individual from both species at the Biomolecular Resource Facility – John Curtin School of Medical Research (BRF-JCSMR) Australian National University (ANU). In short, tissues were fixed in formaldehyde, DNA was extracted and then digested with Arima restriction enzymes. Overhangs of digested DNA were then end repaired and marked with biotinylated nucleotides. The ends were then ligated together, and DNA strands were de-crosslinked and sheared for Illumina PE 150bp sequencing. The *E. fornicatus* library was sequenced on an Illumina NovaSeq 6000 for 300 cycles and the library for *E. similis* was sequenced on an Illumina NextSeq 2000 for 300 cycles.

The Hi-C reads were mapped to draft contig assemblies following the Arima Genomics Mapping Pipeline v03. First, 5’ ends of the Hi-C reads were trimmed by 5 base pairs to remove over-represented 3bp barcode UMI sequences and 2 dark bases from the 5’ end. Trimmed reads were then mapped independently (as single-ended reads) using BWA-MEM2 (Vasimuddin et al. 2019) with the -5SP flag. Only the 5’ ends were retained using the “filter_five_end.pl” script provided by the Arima Genomics Mapping Pipeline, as the ends of the DNA were ligated together prior to sequencing, 3’ ends can originate from the same contiguous strands of DNA as the 5’ end of the mate-pair, which can manifest a ligation function and form chimeric reads after mapping. Following this, single-ended mapped reads were then paired using the “two_read_bam_combiner.pl” script provided by the Arima Genomics Mapping Pipeline and mapped reads with a mapping quality (MAPQ) < 10 were discarded and sorted with samtools (Danecek et al. 2021). Lastly, PCR-duplicates were removed with Picard Tools v3. Assemblies were then scaffolded using YahS v1.1a (Zhou et al. 2023) and assessed with QUAST v5.2.0, and BUSCO v5.5.0 (Manni et al. 2021). We then compared the best scaffolded assembly with an analogous scaffolding run using SALSA2 v2.3 (Ghurye et al. 2019) and selected the best scaffolded assembly using QUAST and BUSCO metrics for downstream curation with the Juicebox Assembly Tools (JBAT) v1.b (Durand et al. 2016). Contaminant scaffolds, defined as contigs identified not to be of Arthropoda origin through BLAST, sequences with >15x coverage, and GC <0.3 and >0.45 were removed using the BlobToolKit v4.2.1 (Challis et al. 2020). Hi-C reads were then mapped to the decontaminated scaffolds using BWA-MEM2 following the Arima Genomics Mapping Pipeline v03 and were reviewed with JBAT v1.b (Durand et al. 2016). Illumina short read and PacBio HiFi reads were also mapped to the decontaminated scaffolds and all three mapped datasets were inspected using JBrowse v2.10.3 to identify any potential misassembles (identified by poor read alignments and regions of low coverage). We assessed the completeness of the final scaffolded assemblies again with QUAST v5.2.0 (Mikheenko et al. 2018) and BUSCO v5.5.0 (Manni et al. 2021). Lastly, the scaffolds of each library of the same species (i.e. both *E. fornicatus* and *E. similis* assemblies) were renamed and reoriented to be syntenic with one another using BUSCOs to define synteny and orientation. We identified pseudo-chromosome scaffolds by scaffolds that were > 500Kb in length and had an average Hi-C coverage > 30x. Pairs of syntenic pseudo-chromosomes were defined by length and by BUSCO composition and these pseudo-chromosomes were named in order of length by the EFF26 assembly for *E. fornicatus* and the ESF13 assembly for *E. similis*. Lastly, assemblies were assessed for any remaining contamination with FCS-GX v0.5 (Astashyn et al. 2024) and any contaminants identified were removed prior to submission to the NCBI GenBank.

### Genome annotation

Total RNA was extracted from ten adult males and females of each species. RNA was extracted using the Zymo Research Quick-RNA Miniprep Plus kit following the manufacturers tough-to-lyse sample preparation procedure. RNA quantity and quality was then assessed using a DeNovix spectrophotometer and Aligent Fragment Analyzer system with the HS RNA kit DNF-472. For one extract of each species and sex strand-specific libraries were prepared by GENEWIZ from Azenta Life Sciences and sequenced on an Illumina NovaSeq 6000 PE150.

Genomes were annotated by The NCBI Eukaryotic Annotation Pipeline. In short, the NCBI GenBank reference assemblies of *E. fornicatus* (EFF26) and *E. similis* (ESF13) were masked with WindowMasker (Morgulis et al. 2006), and available transcripts, RNA-seq (which includes data produced herein), and protein data from RefSeq were mapped to masked genomes. Transcripts and proteins from the RefSeq and GenBank databases were aligned with BLAST and the short-read RNA-seq data was aligned with STAR (Dobin et al. 2013). The final set of genes were built by evaluation against known RefSeq transcripts, curated RefSeq genomic alignments and the most highly supported models that were predicted by Gnomon. Protein naming, locus type and GeneID assignment were made by the NCBI Eukaryotic Annotation Pipeline. Completeness of the annotated gene models was then assessed with BUSCO v5.5.0 (Manni et al. 2021) run in protein mode using the Endopterygota_obd10 dataset and the tribolium2012 dataset was specified for Augustus (Stanke et al. 2008) training. The distribution and density of genes along chromosomes of EFF26 and ESF13 were visualised with RIdeogram (Hao et al. 2020).

### Transposable Elements

Chromosome level genomes of *Taphrorychus bicolor* (Telfer et al. 2024), *Dendroctonus ponderosae* (Keeling et al. 2022)*, D. valens* (Liu et al. 2022)*, Ips nitidus* (Wang et al. 2023) and the outgroup *Polydrsus cervinus* (Entiminae) (Barclay et al. 2023) were downloaded from NCBI or ensemble. First, repeat annotation was performed with EarlGreyTE (Baril et al. 2024) to identify, classify and mask repeat regions across published genomes and the reference assemblies of *E. fornicatus* (EFF26) and *E. similis* (ESF13). Following repeat annotation, kimura-2 divergences between TE classes were calculated using the parseRM.pl scrip with the -l flag (Kapusta 2017).

The composition of TEs were compared phylogenetically across Scolytinae genomes downloaded and the *Euwallacea* reference genomes (EFF26 & ESF13). We used BUSCO to call 2,108 single copy sequences which were aligned with MAFFT v7.490 (Katoh 2002) and poorly aligned bases were removed with TrimAl v1.4.1r22 (Capella-Gutiérrez et al. 2009). Alignments were then concatenated with phykit v1.19.0 (Steenwyk et al. 2021) and phylogenetic analyses were conducted in IQ-tree v2.3.2 (Minh et al. 2020). Substitution models were estimated with ModelFinder Plus (Kalyaanamoorthy et al. 2017) using a greedy strategy with gene merging and bootstrap support was calculated with nonparametric bootstrap with 100. The relative proportions of TEs were then annotated on the tips of the phylogeny.

### Chromosomal synteny across Scolytinae

Macrosynteny and chromosomal variation across published Scolytinae genomes and the *Euwallacea* reference genomes (EFF26 & ESF13) was performed with the Oxford Dot Plots (opd) pipeline (Schultz et al. 2023). Previously used BUSCO sequences were used to describe the relationship between linkage groups and chromosomes between species. Reciprocal best hits between BUSCOs were identified with diamond blastp and the homology of scaffolds between species were inferred with pairwise p-values calculated with Fisher’s exact test. Pairwise species dot plots syntenic linkage ribbon plots for all species were generated using scripts available in odp. Linkage colours in the ribbon plot follows the Bilaterian-Cnidarian-Sponge Linkage Groups (BCnS LGs) as in (Schultz et al. 2023).

### Mitochondrial phylogenetics

To identify the haplotype of *E. fornicatus* present in Perth WA, previously published *COI* genes (Stouthamer et al. 2017; Gomez et al. 2018; Bierman et al. 2022) of PSHB, TSHBa&b and KSHB were downloaded from NCBI. The mitochondrial genome for all libraries were assembled using the MitoHiFi v3.2.1 pipeline (Uliano-Silva et al. 2023). Full length *COI* genes from these assemblies were then aligned with the sequences from NCBI using MAFFT (Katoh 2002). Third codon positions were partitioned in the alignment and IQ-Tree v2.3.2 (Minh et al. 2020) was used to estimate the phylogeny, substitution models were identified with ModelFinderPlus (Kalyaanamoorthy et al. 2017) and bootstrap support was calculated with UFBoot (Hoang et al. 2018) with 1000 replicates.

### Metagenomics

To identify the microbial communities associated with *E. fornicatus* and *E. similis*, metagenomes were assembled from the total hifi sequence read dataset for each library. Metagenome-assembled genomes (MAGs) were assembled using hifiasm-meta (Feng et al. 2022) with default settings. BLAST was used to identify contigs and MAG identity was explored and visualised with the BlobToolKit v4.2.1 (Challis et al. 2020). Host contigs and those that were unidentified were filtered out of the dataset. A closed *Wolbachia* contig in was found in both *E. similis* libraries, and so the strain was identified using MLST approaches implemented in the PubMLST database (Jolley et al. 2018).

## Data Availability

All genome and raw sequencing reads produced in this have been deposited with the NCBI, under the BioProject: PRJNA1086994. Specific accession numbers for raw sequence reads and genome assemblies are available in Table S1 and Table 1, respectively. Vouchers of the specimens are deposited in the ANIC, CSIRO under accession numbers: ANIC-25-086063 – ANIC-25-086073.

## Author’s Contributions

JB, HE, TW, RR, VC, AS and TW conceptualised the research. DPIRD and KI provided PSHB specimens. TW, RR, LC and GP provided advice for HWM DNA extraction and genome assembly pipelines. JB and HE interpreted the results. JB conducted the field and laboratory work, outlined the bioinformatic pipelines, collected and analysed the data, and wrote the manuscript with input from all authors. All authors agree to the submitted manuscript.

## Acknowledgements

We are grateful to the Polyphagous Shot Hole Borer response team from the Department of Primary Industries Regional Development Western Australia for providing samples, including Basman Al-Jalely and Emad Alshwany from the Reproductive Biology Laboratory. Bonnie Koopmans and Debbie Jennings for their help during fieldwork. We would like to thank Andreas Zwick, James Nicholls, Diana Hartley and Andrew Gock (CSIRO Black Mountain) for access to laboratory facilities, Peter Holland (Oxford University, UK) and Mara Lawniczak (Darwin Tree of Life, UK) for advice on laboratory protocols. We thank Maxim Nekrasov and Carolina Correa Ospina (Australian National University Biomolecular Resource Facility), and Trent Peters and Oliver Distler (Australian Genome Research Facility), for their technical support. The authors thank John G. Oakeshott (CSIRO) and Robert Waterhouse (University of Lausanne) for their generous advice and David Yeates (CSIRO Black Mountain) for his support of the project. We would like to thank Andy Bachler for advice on genome scaffolding and Ondrej Hlinka (CSIRO) for assistance with High Performance Computing. We also thank Cecile Gueidan (CSIRO) for comments on an earlier version of this manuscript. Thanks to CSIRO research Office and the Zimmerman Trust for supporting this research.

## References

Astashyn A, et al. 2024. Rapid and sensitive detection of genome contamination at scale with FCS-GX. Genome Biol. 25(60). 10.1186/s13059-024-03198-7.

Barclay MVL, et al. 2023. The genome sequence of a weevil, *Polydrusus cervinus* (Linnaeus, 1758). Wellcome Open Res. 8(563). 10.12688/wellcomeopenres.20414.1.

Baril T, Galbraith J, Hayward A. 2024. Earl Grey: A fully automated user-friendly transposable element annotation and analysis pipeline. Molecular Biology and Evolution. 41(4):msae068. 10.1093/molbev/msae068.

Beaver RA, Sittichaya W, Liu L-Y. 2014. A Synopsis of the scolytine ambrosia beetles of Thailand (Coleoptera: Curculionidae: Scolytinae). Zootaxa. 3875(1):1–82. 10.11646/zootaxa.3875.1.1.

Bickerstaff JRM, Jordal BH, Riegler M. 2023. Two sympatric lineages of Australian *Cnestus solidus* share *Ambrosiella* symbionts but not *Wolbachia*. Heredity. 132(1):43–53. 10.1038/s41437-023-00659-w.

Biedermann PHW, et al. 2019. Bark beetle population dynamics in the Anthropocene: Challenges and solutions. Trends in Ecology & Evolution. 34(1):914–924. 10.1016/j.tree.2019.06.002.

Bierman A, Roets F, Terblanche JS. 2022. Population structure of the invasive ambrosia beetle, *Euwallacea fornicatus*, indicates multiple introductions into South Africa. Biol Invasions. 24(8):2301–2312. 10.1007/s10530-022-02801-x.

Blackmon H, Demuth JP. 2015. Coleoptera karyotype database. The Coleopterists Bulletin. 69(1):174–175. 10.1649/0010-065X-69.1.174.

Blackmon H, Jonika MM, Alfieri JM, Fardoun L, Demuth JP. 2024. Drift drives the evolution of chromosome number I: The impact of trait transitions on genome evolution in Coleoptera. Journal of Heredity. 115(2):173–182. 10.1093/jhered/esae001.

Capella-Gutiérrez S, Silla-Martínez JM, Gabaldón T. 2009. trimAl: a tool for automated alignment trimming in large-scale phylogenetic analyses. Bioinformatics. 25(15):1972–1973. 10.1093/bioinformatics/btp348.

Challis R, Richards E, Rajan J, Cochrane G, Blaxter M. 2020. BlobToolKit – Interactive quality assessment of genome assemblies. G3 Genes|Genomes|Genetics. 10(4):1361–1374. 10.1534/g3.119.400908.

Cheng H, Concepcion GT, Feng X, Zhang H, Li H. 2021. Haplotype-resolved de novo assembly using phased assembly graphs with hifiasm. Nat Methods. 18(2):170–175. 10.1038/s41592-020-01056-5.

Cook DC, Broughton S. 2023. Economic impact of polyphagous shot hole borer *Euwallacea fornicatus* (Coleoptera: Curculionidae: Scolytinae) in Western Australia. Agri and Forest Entomology. 25(3):449–457. 10.1111/afe.12566.

Danecek P, et al. 2021. Twelve years of SAMtools and BCFtools. GigaScience. 10(2):giab008. 10.1093/gigascience/giab008.

De Jager MM, Roets F. 2022. Rapid and cost-effective detection of *Fusarium euwallaceae* from woody tissues. Plant Pathology. 71(8):1712–1720. 10.1111/ppa.13600.

De Wit MP, et al. 2022. An Assessment of the Potential Economic Impacts of the Invasive Polyphagous Shot Hole Borer (Coleoptera: Curculionidae) in South Africa. Journal of Economic Entomology. 115(4):1076–1086. 10.1093/jee/toac061.

Dobin A, et al. 2013. STAR: ultrafast universal RNA-seq aligner. Bioinformatics. 29(1):15–21. 10.1093/bioinformatics/bts635.

Durand NC, et al. 2016. Juicebox provides a visualization system for Hi-C contact maps with unlimited zoom. Cell Systems. 3(1):99–101. 10.1016/j.cels.2015.07.012.

Eskalen A, et al. 2012. First report of a *Fusarium* sp. and its vector Tea Shot Hole Borer (*Euwallacea fornicatus*) causing *Fusarium* dieback on avocado in California. Plant Disease. 96(7):1070–1070. 10.1094/PDIS-03-12-0276-PDN.

Feng X, Cheng H, Portik D, Li H. 2022. Metagenome assembly of high-fidelity long reads with hifiasm-meta. Nat Methods. 19(6):671–674. 10.1038/s41592-022-01478-3.

Feschotte C, Pritham EJ. 2007. DNA transposons and the evolution of Eukaryotic genomes. Annu. Rev. Genet. 41(1):331–368. 10.1146/annurev.genet.40.110405.090448.

Freeman S, et al. 2013. *Fusarium euwallaceae* sp. nov.—a symbiotic fungus of *Euwallacea* sp., an invasive ambrosia beetle in Israel and California. Mycologia. 105(6):1595–1606. 10.3852/13-066.

Frey JE, et al. 2022. Next generation biosecurity: Towards genome based identification to prevent spread of agronomic pests and pathogens using nanopore sequencing. PLoS ONE. 17(7):e0270897. 10.1371/journal.pone.0270897.

Gerdol M, et al. 2020. Massive gene presence-absence variation shapes an open pan-genome in the Mediterranean mussel. Genome Biol. 21(1):275. 10.1186/s13059-020-02180-3.

Ghurye J, et al. 2019. Integrating Hi-C links with assembly graphs for chromosome-scale assembly. PLoS Comput Biol. 15(8):e1007273. 10.1371/journal.pcbi.1007273.

Gilbert C, Peccoud J, Cordaux R. 2021. Transposable elements and the evolution of insects. Annu. Rev. Entomol. 66(1):355–372. 10.1146/annurev-ento-070720-074650.

Gomez DF, et al. 2018. Species delineation within the *Euwallacea fornicatus* (Coleoptera: Curculionidae) complex revealed by morphometric and phylogenetic analyses. Insect Systematics and Diversity. 2:(6)2–11. 10.1093/isd/ixy018.

Gomez DF, Lin W, Gao L, Li Y. 2019. New host plant records for the *Euwallacea fornicatus* (Eichhoff) species complex (Coleoptera: Curculionidae: Scolytinae) across its natural and introduced distribution. Journal of Asia-Pacific Entomology. 22(1):338–340. 10.1016/j.aspen.2019.01.013.

Guan D, et al. 2020. Identifying and removing haplotypic duplication in primary genome assemblies. Bioinformatics. 36(9):2896–2898. 10.1093/bioinformatics/btaa025.

Guo Q, Huang W, Sang W, Chen X, Wang X. 2023. Characterization, comparative analyses, and phylogenetic implications of mitochondrial genomes among bark and ambrosia beetles (Coleoptera: Curculionidae, Scolytinae). Front. Ecol. Evol. 11:1191446. 10.3389/fevo.2023.1191446.

Hao Z, et al. 2020. *RIdeogram*: drawing SVG graphics to visualize and map genome-wide data on the idiograms. PeerJ Computer Science. 6:e251. 10.7717/peerj-cs.251.

Hoang DT, Chernomor O, Von Haeseler A, Minh BQ, Vinh LS. 2018. UFBoot2: Improving the ultrafast bootstrap approximation. Molecular Biology and Evolution. 35(2):518–522. 10.1093/molbev/msx281.

Hulcr J, Stelinski LL. 2017. The ambrosia symbiosis: From evolutionary ecology to practical management. Annu. Rev. Entomol. 62(1):285–303. 10.1146/annurev-ento-031616-035105.

Jolley KA, Bray JE, Maiden MCJ. 2018. Open-access bacterial population genomics: BIGSdb software, the PubMLST.org website and their applications. Wellcome Open Res. 3:124. 10.12688/wellcomeopenres.14826.1.

Jordal B. 2014. 3.7. 12 Scolytinae Latreille, 1806. Coleoptera, Beetles. Volume 3: Morphology and Systematics (Phytophaga). 633–642. Walter de Gruyter Berlin and New York.

Jordal BH, Beaver RA, Kirkendall LR. 2001. Breaking taboos in the tropics: incest promotes colonization by wood-boring beetles. Global Ecology and Biogeography. 10(4):345–357. 10.1046/j.1466-822X.2001.00242.x.

Jordal BH, Cognato AI. 2012. Molecular phylogeny of bark and ambrosia beetles reveals multiple origins of fungus farming during periods of global warming. BMC Evol Biol. 12(1):133. 10.1186/1471-2148-12-133.

Kalyaanamoorthy S, Minh BQ, Wong TKF, Von Haeseler A, Jermiin LS. 2017. ModelFinder: fast model selection for accurate phylogenetic estimates. Nat Methods. 14(6):587–589. 10.1038/nmeth.4285.

Kapusta. 2017. Parsing-RepeatMasker-Outputs. https://github.com/4ureliek/Parsing-RepeatMasker-Outputs.

Katoh K. 2002. MAFFT: a novel method for rapid multiple sequence alignment based on fast Fourier transform. Nucleic Acids Research. 30(14):3059–3066. 10.1093/nar/gkf436.

Kawasaki Y, Schuler H, Stauffer C, Lakatos F, Kajimura H. 2016. *Wolbachia* endosymbionts in haplodiploid and diploid scolytine beetles (Coleoptera: Curculionidae: Scolytinae). Environ Microbiol Rep. 8(5):680–688. 10.1111/1758-2229.12425.

Keeling CI, et al. 2022. Chromosome-level genome assembly reveals genomic architecture of northern range expansion in the mountain pine beetle, *Dendroctonus ponderosae* Hopkins (Coleoptera: Curculionidae). Molecular Ecology Resources. 22(3):1149–1167. 10.1111/1755-0998.13528.

Lanier GN. 1981. Cytotaxonomy of *Dendroctonus*. Forest, Wildlife, and Range Experiment Station, University of Idaho, Moscow.

Lanier GN, Kirkendall LR. 1986. Karyology of pseudogamous *Ips* bark beetles. Hereditas. 105(1):87–96. doi: 10.1111/j.1601-5223.1986.tb00646.x.

Lantschner MV, Corley JC, Liebhold AM. 2020. Drivers of global Scolytinae invasion patterns. Ecological Applications. 30(5):e02103. 10.1002/eap.2103.

Lester PJ, et al. 2020. The potential for a CRISPR gene drive to eradicate or suppress globally invasive social wasps. Sci Rep. 10(1):12398. 10.1038/s41598-020-69259-6.

Li K, Xu P, Wang J, Yi X, Jiao Y. 2023. Identification of errors in draft genome assemblies at single-nucleotide resolution for quality assessment and improvement. Nat Commun. 14(1):6556. 10.1038/s41467-023-42336-w.

Liu Z et al. 2022. Chromosome-level genome assembly and population genomic analyses provide insights into adaptive evolution of the red turpentine beetle, *Dendroctonus valens*. BMC Biol. 20(1):190. 10.1186/s12915-022-01388-y.

Lynn KMT, et al. 2021. Novel *Fusarium* mutualists of two *Euwallacea* species infesting *Acacia crassicarpa* in Indonesia. Mycologia. 113(3):536–558. 10.1080/00275514.2021.1875708.

Manni M, Berkeley MR, Seppey M, Simão FA, Zdobnov EM. 2021. BUSCO Update: Novel and streamlined workflows along with broader and deeper phylogenetic coverage for scoring of Eukaryotic, Prokaryotic, and Viral genomes. Molecular Biology and Evolution. 38(10):4647–4654. 10.1093/molbev/msab199.

Marçais G, Kingsford C. 2011. A fast, lock-free approach for efficient parallel counting of occurrences of *k* -mers. Bioinformatics. 27(6):764–770. 10.1093/bioinformatics/btr011.

McCutcheon JP, Moran NA. 2012. Extreme genome reduction in symbiotic bacteria. Nat Rev Microbiol. 10(1):13–26. 10.1038/nrmicro2670.

McGaughran A, et al. 2024. Genomic tools in biological invasions: Current state and future frontiers. Genome Biology and Evolution. 16(1):evad230. 10.1093/gbe/evad230.

Meiborg AB, Faber NR, Taylor BA, Harpur BA, Gorjanc G. 2023. The suppressive potential of a gene drive in populations of invasive social wasps is currently limited. Sci Rep. 13(1):1640. 10.1038/s41598-023-28867-8.

Mendel Z, et al. 2012. An Asian ambrosia beetle *Euwallacea fornicatus* and its novel symbiotic fungus *Fusarium* sp. pose a serious threat to the Israeli avocado industry. Phytoparasitica. 40(3):235–238. 10.1007/s12600-012-0223-7.

Mendel Z, et al. 2021. What determines host range and reproductive performance of an invasive ambrosia beetle *Euwallacea fornicatus*; Lessons from Israel and California. Front. For. Glob. Change. 4:654702. 10.3389/ffgc.2021.654702.

Mikheenko A, Prjibelski A, Saveliev V, Antipov D, Gurevich A. 2018. Versatile genome assembly evaluation with QUAST-LG. Bioinformatics. 34(13):i142–i150. 10.1093/bioinformatics/bty266.

Minh BQ, et al. 2020. IQ-TREE 2: New models and efficient methods for phylogenetic inference in the genomic era. Molecular Biology and Evolution. 37(5):1530–1534. 10.1093/molbev/msaa015.

Moggioli G, et al. 2023. Distinct genomic routes underlie transitions to specialised symbiotic lifestyles in deep-sea annelid worms. Nat Commun. 14(1):2814. 10.1038/s41467-023-38521-6.

Morgulis A, Gertz EM, Schäffer AA, Agarwala R. 2006. WindowMasker: window-based masker for sequenced genomes. Bioinformatics. 22(2):134–141. 10.1093/bioinformatics/bti774.

Navarro-Escalante L, et al. 2021. A coffee berry borer (*Hypothenemus hampei*) genome assembly reveals a reduced chemosensory receptor gene repertoire and male-specific genome sequences. Sci Rep. 11(1):4900. 10.1038/s41598-021-84068-1.

North HL, McGaughran A, Jiggins CD. 2021. Insights into invasive species from whole-genome resequencing. Molecular Ecology. 30(23):6289–6308. 10.1111/mec.15999.

Nurk S, et al. 2020. HiCanu: accurate assembly of segmental duplications, satellites, and allelic variants from high-fidelity long reads. Genome Res. 30(9):1291–1305. 10.1101/gr.263566.120.

Nygaard S, et al. 2011. The genome of the leaf-cutting ant *Acromyrmex echinatior* suggests key adaptations to advanced social life and fungus farming. Genome Res. 21(8):1339–1348. 10.1101/gr.121392.111.

Nygaard S, et al. 2016. Reciprocal genomic evolution in the ant–fungus agricultural symbiosis. Nat Commun. 7(1):12233. 10.1038/ncomms12233.

O’Donnell K, et al. 2016. Invasive Asian *Fusarium* – *Euwallacea* ambrosia beetle mutualists pose a serious threat to forests, urban landscapes and the avocado industry. Phytoparasitica. 44(4):435–442. 10.1007/s12600-016-0543-0.

Paap T, De Beer ZW, Migliorini D, Nel WJ, Wingfield MJ. 2018. The polyphagous shot hole borer (PSHB) and its fungal symbiont *Fusarium euwallaceae*: a new invasion in South Africa. Australasian Plant Pathol. 47(2):231–237. 10.1007/s13313-018-0545-0.

Peer K, Taborsky M. 2005. Outbreeding depression, but no inbreeding depression in haplodiploid ambrosia beetles with regular sibling mating. Evolution. 59(2):317–323. 10.1111/j.0014-3820.2005.tb00992.x.

Petersen M, et al. 2019. Diversity and evolution of the transposable element repertoire in arthropods with particular reference to insects. BMC Evol Biol. 19(1):11. 10.1186/s12862-018-1324-9.

Pistone D, Gohli J, Jordal BH. 2018. Molecular phylogeny of bark and ambrosia beetles (Curculionidae: Scolytinae) based on 18 molecular markers. Systematic Entomology. 43(2):387–406. 10.1111/syen.12281.

Powell D, et al. 2021. A highly-contiguous genome assembly of the Eurasian spruce bark beetle, Ips typographus, provides insight into a major forest pest. Commun Biol. 4(1):1059. 10.1038/s42003-021-02602-3.

Ranallo-Benavidez TR, Jaron KS, Schatz MC. 2020. GenomeScope 2.0 and Smudgeplot for reference-free profiling of polyploid genomes. Nat Commun. 11(1):1432. 10.1038/s41467-020-14998-3.

Rane R et al. 2023. Complex multiple introductions drive fall armyworm invasions into Asia and Australia. Sci Rep. 13(1):660. 10.1038/s41598-023-27501-x.

Ren L, Huang W, Cannon EKS, Bertioli DJ, Cannon SB. 2018. A mechanism for genome size reduction following genomic rearrangements. Front. Genet. 9:454. 10.3389/fgene.2018.00454.

Ross PA, et al. 2022. A decade of stability for *w*Mel Wolbachia in natural *Aedes aegypti* populations. PLoS Pathog. 18(2):e1010256. 10.1371/journal.ppat.1010256.

Rugman-Jones PF, et al. 2020. One becomes two: second species of the *Euwallacea fornicatus* (Coleoptera: Curculionidae: Scolytinae) species complex is established on two Hawaiian Islands. PeerJ. 8:e9987. 10.7717/peerj.9987.

Ruzzier E, et al. 2023. The first full host plant dataset of Curculionidae Scolytinae of the world: tribe Xyleborini LeConte, 1876. Sci Data. 10(1):166. 10.1038/s41597-023-02083-5.

Schrader L, et al. 2021. Relaxed selection underlies genome erosion in socially parasitic ant species. Nat Commun. 12(1):2918. 10.1038/s41467-021-23178-w.

Schuler H, et al. 2023. Recent invasion and eradication of two members of the *Euwallacea fornicatus* species complex (Coleoptera: Curculionidae: Scolytinae) from tropical greenhouses in Europe. Biol Invasions. 25(2):299–307. 10.1007/s10530-022-02929-w.

Schultz DT, et al. 2023. Ancient gene linkages support ctenophores as sister to other animals. Nature. 618(7963):110–117. 10.1038/s41586-023-05936-6.

Sillero N, Huey RB, Gilchrist G, Rissler L, Pascual M. 2020. Distribution modelling of an introduced species: do adaptive genetic markers affect potential range? Proc. R. Soc. B. 287(1935):20201791. 10.1098/rspb.2020.1791.

Sim SB, Corpuz RL, Simmonds TJ, Geib SM. 2022. HiFiAdapterFilt, a memory efficient read processing pipeline, prevents occurrence of adapter sequence in PacBio HiFi reads and their negative impacts on genome assembly. BMC Genomics. 23(1):157. 10.1186/s12864-022-08375-1.

Six DL, Bracewell R. 2015. *Dendroctonus*. In: Bark Beetles. Elsevier pp. 305–350. 10.1016/B978-0-12-417156-5.00008-3.

Sloan DB, Moran NA. 2012. Genome reduction and co-evolution between the primary and secondary bacterial symbionts of psyllids. Molecular Biology and Evolution. 29(12):3781–3792. 10.1093/molbev/mss180.

Smith SM, F. Gomez D, A. Beaver R, Hulcr J, I. Cognato A. 2019. Reassessment of the species in the *Euwallacea fornicatus* (Coleoptera: Curculionidae: Scolytinae) complex after the rediscovery of the “lost” type specimen. Insects. 10(9):261. 10.3390/insects10090261.

Stanke M, Diekhans M, Baertsch R, Haussler D. 2008. Using native and syntenically mapped cDNA alignments to improve *de novo* gene finding. Bioinformatics. 24:637–644. doi: 10.1093/bioinformatics/btn013.

Stapley J, Santure AW, Dennis SR. 2015. Transposable elements as agents of rapid adaptation may explain the genetic paradox of invasive species. Molecular Ecology. 24(9):2241–2252. 10.1111/mec.13089.

Steenwyk JL et al. 2021. PhyKIT: a broadly applicable UNIX shell toolkit for processing and analyzing phylogenomic data. Bioinformatics. 37(16):2325–2331. 10.1093/bioinformatics/btab096.

Storer CG, Breinholt JW, Hulcr J. 2015. *Wallacellus* is *Euwallacea*: molecular phylogenetics settles generic relationships (Coleoptera: Curculionidae: Scolytinae: Xyleborini). Zootaxa. 3974(3):391–400. 10.11646/zootaxa.3974.3.6.

Stouthamer R, et al. 2017. Tracing the origin of a cryptic invader: phylogeography of the *Euwallacea fornicatus* (Coleoptera: Curculionidae: Scolytinae) species complex. Agri and Forest Entomology. 19(4):366–375. 10.1111/afe.12215.

Takenouchi Y, Takagi K. 1967. A chromosome study of two parthenogenetic scolytid beetles. Annotationes zoologicae japonenses. 40(2): 105–110.

Telfer MG, et al. 2024. The genome sequence of the beech bark beetle, *Taphrorychus bicolor* (Herbst, 1793). Wellcome Open Res. 9:213. 10.12688/wellcomeopenres.21265.1.

Uliano-Silva M, et al. 2023. MitoHiFi: a python pipeline for mitochondrial genome assembly from PacBio high fidelity reads. BMC Bioinformatics. 24(1):288. 10.1186/s12859-023-05385-y.

Vasimuddin Md, Misra S, Li H, Aluru S. 2019. Efficient architecture-aware acceleration of BWA-MEM for multicore systems. In: 2019 IEEE International Parallel and Distributed Processing Symposium (IPDPS). pp. 314–324. 10.1109/IPDPS.2019.00041.

Vilardo G, Faccoli M, Corley JC, Lantschner MV. 2022. Factors driving historic intercontinental invasions of European pine bark beetles. Biol Invasions. 24(9):2973–2991. 10.1007/s10530-022-02818-2.

Virkki N, Smith SG. 1978. *Animal Cytogenetics*. Schweizerbart Science Publishers: Stuttgart, Germany

Wang L-J, Hsu M-H, Liu T-Y, Lin M-Y, Sung C-H. 2020. Characterization of the complete mitochondrial genome of *Euwallacea fornicatus* (Eichhoff, 1868) (Coleoptera: Curculionidae: Scolytinae) and its phylogenetic implications. Mitochondrial DNA Part B. 5(3):3502–3504. 10.1080/23802359.2020.1827070.

Wang Z, et al. 2023. Genome and transcriptome of *Ips nitidus* provide insights into high-altitude hypoxia adaptation and symbiosis. iScience. 26(10):107793. 10.1016/j.isci.2023.107793.

Xi Z, Dean JL, Khoo C, Dobson SL. 2005. Generation of a novel *Wolbachia* infection in *Aedes albopictus* (Asian tiger mosquito) via embryonic microinjection. Insect Biochemistry and Molecular Biology. 35(8):903–910. 10.1016/j.ibmb.2005.03.015.

Zhou C, McCarthy SA, Durbin R. 2023. YaHS: yet another Hi-C scaffolding tool. Bioinformatics. 39(1):btac808. 10.1093/bioinformatics/btac808.

