## Supplementary information for "Chromosome structural rearrangements in invasive haplodiploid ambrosia beetles revealed by the genomes of *Euwallacea fornicatus* and *Euwallacea similis* (Coleoptera, Curculionidae, Scolytinae)"

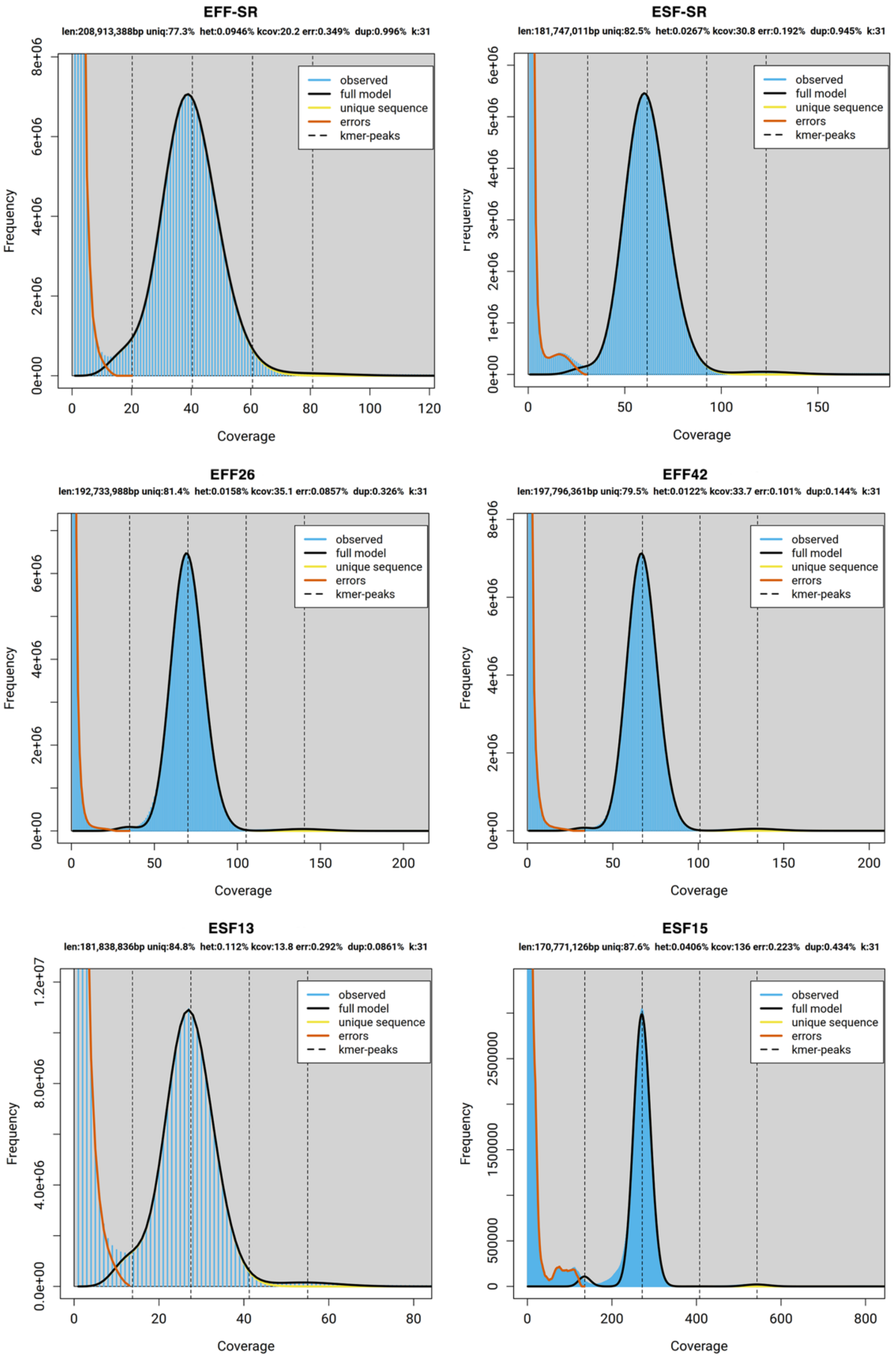


Supplementary Figure 1. GenomeScope (Ranallo-Benavidez et al. 2020; Vurture et al. 2017) profiles generated for Illumina short reads (top row) and PacBio HiFi reads (middle and bottom row), with kmers counted by jellyfish (Marçais & Kingsford 2011). The top left panel is *Euwallacea fornicatus* and the top right panel is *E. similis.* The middle panels are the *Euwallacea fornicatus* long read libraries and the *E. similis* long read libraries are on the bottom.

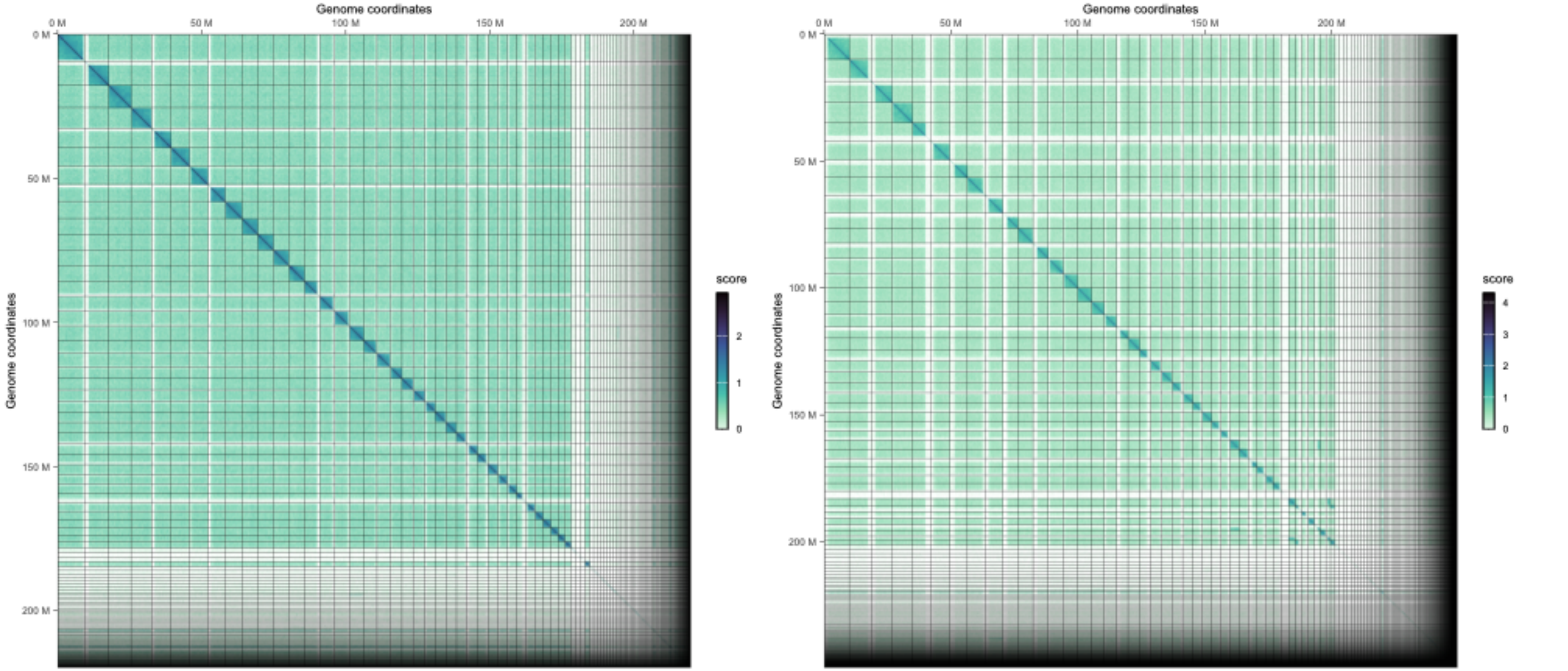


Supplmentary Figure 2. EFF26 contact map of Hi-C reads mapped to the scaffolds generated by YahS (left) (Zhou et al. 2023) and SALSA2 (right) (Ghurye et al. 2019). Contact map was generated using the HiContacts (Serizay et al. 2024) package in R (R Core Team 2013). The Hi-C interaction matrix plotted was coverage normalised and the scale represents a log10 adjusted range of interaction scores.

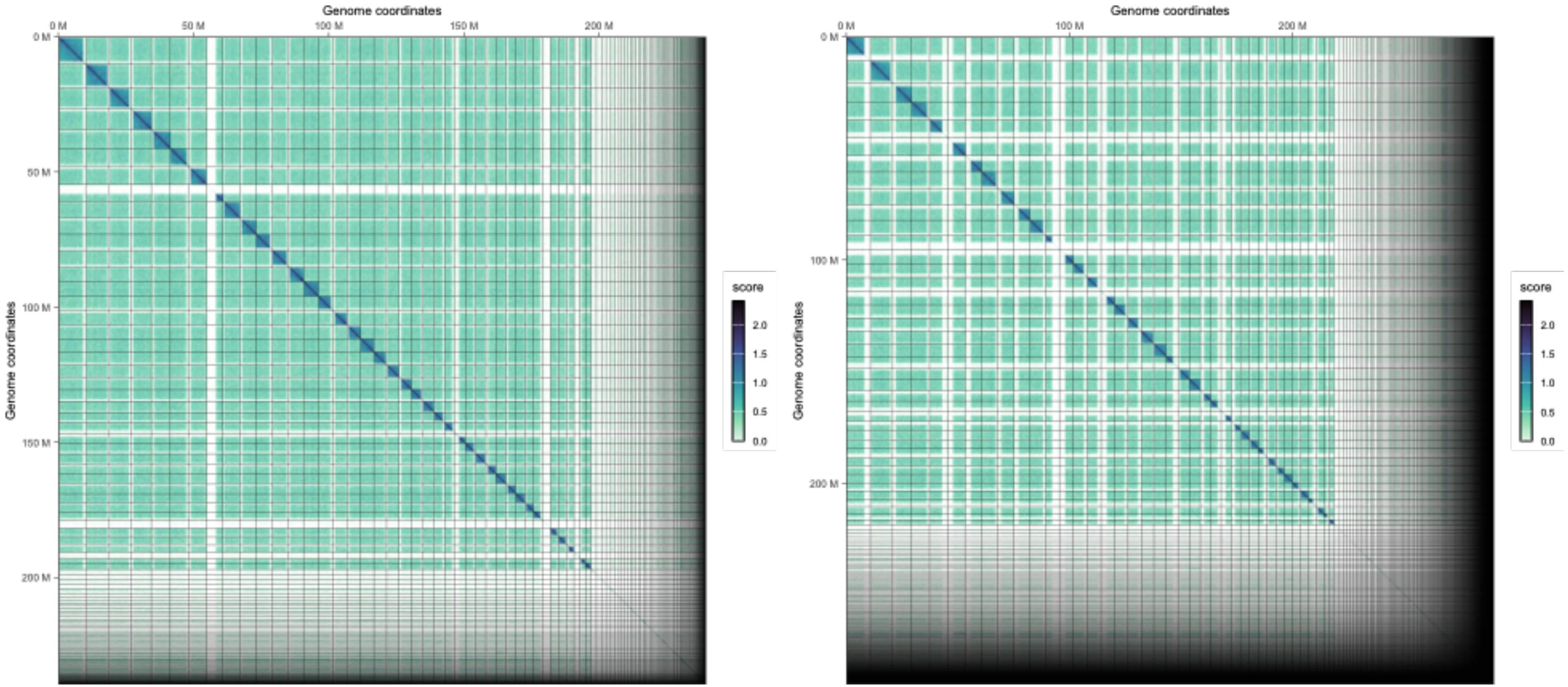


Supplmentary Figure 3. EFF42 contact map of Hi-C reads mapped to the scaffolds generated by YahS (left) (Zhou et al. 2023) and SALSA2 (right) (Ghurye et al. 2019). Contact map was generated using the HiContacts (Serizay et al. 2024) package in R (R Core Team 2013). The Hi-C interaction matrix plotted was coverage normalised and the scale represents a log10 adjusted range of interaction scores.

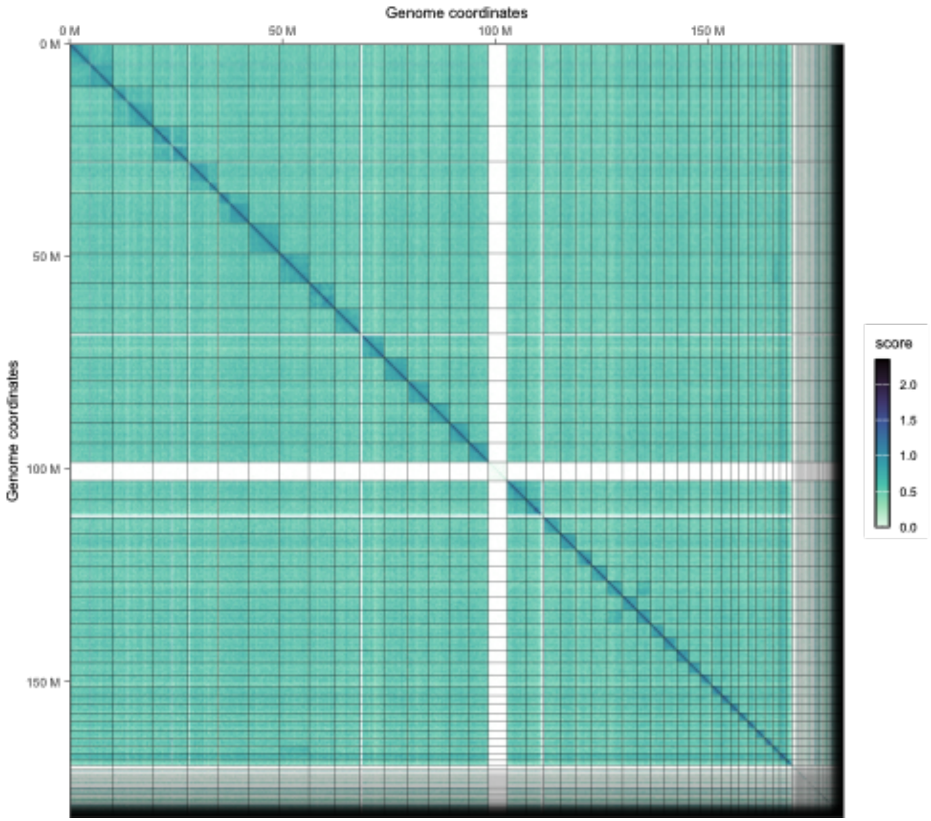


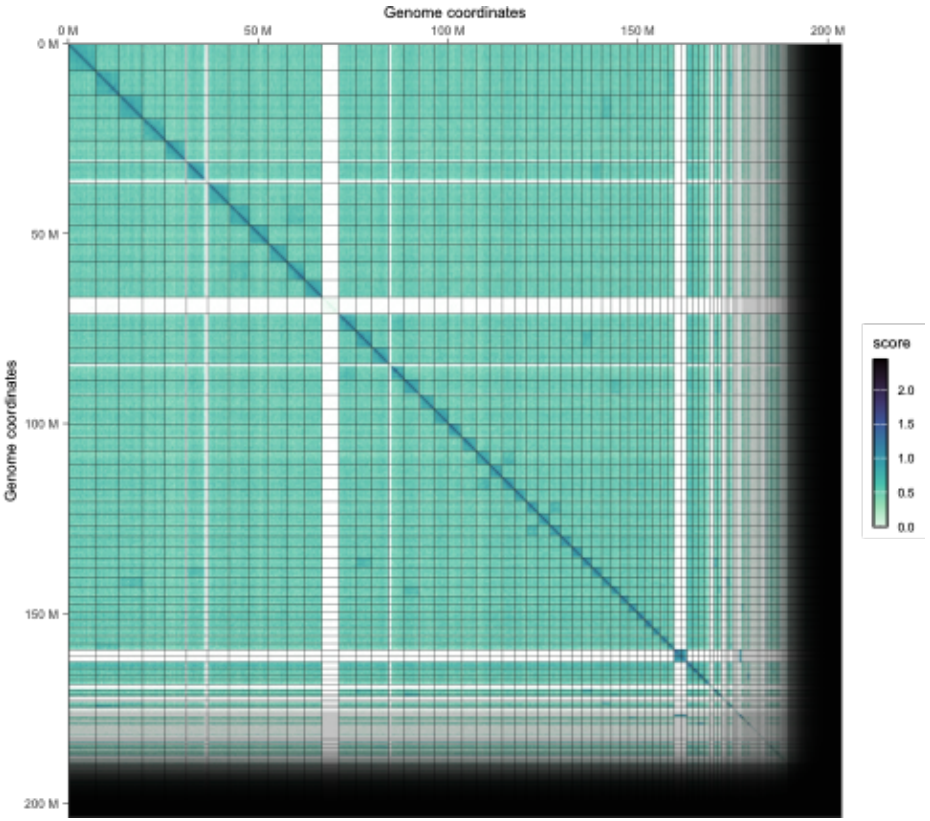


Supplmentary Figure 4. ESF13 contact map of Hi-C reads mapped to the scaffolds generated by YahS (left) (Zhou et al. 2023) and SALSA2 (right) (Ghurye et al. 2019). Contact map was generated using the HiContacts (Serizay et al. 2024) package in R (R Core Team 2013). The Hi-C interaction matrix plotted was coverage normalised and the scale represents a log10 adjusted range of interaction scores.

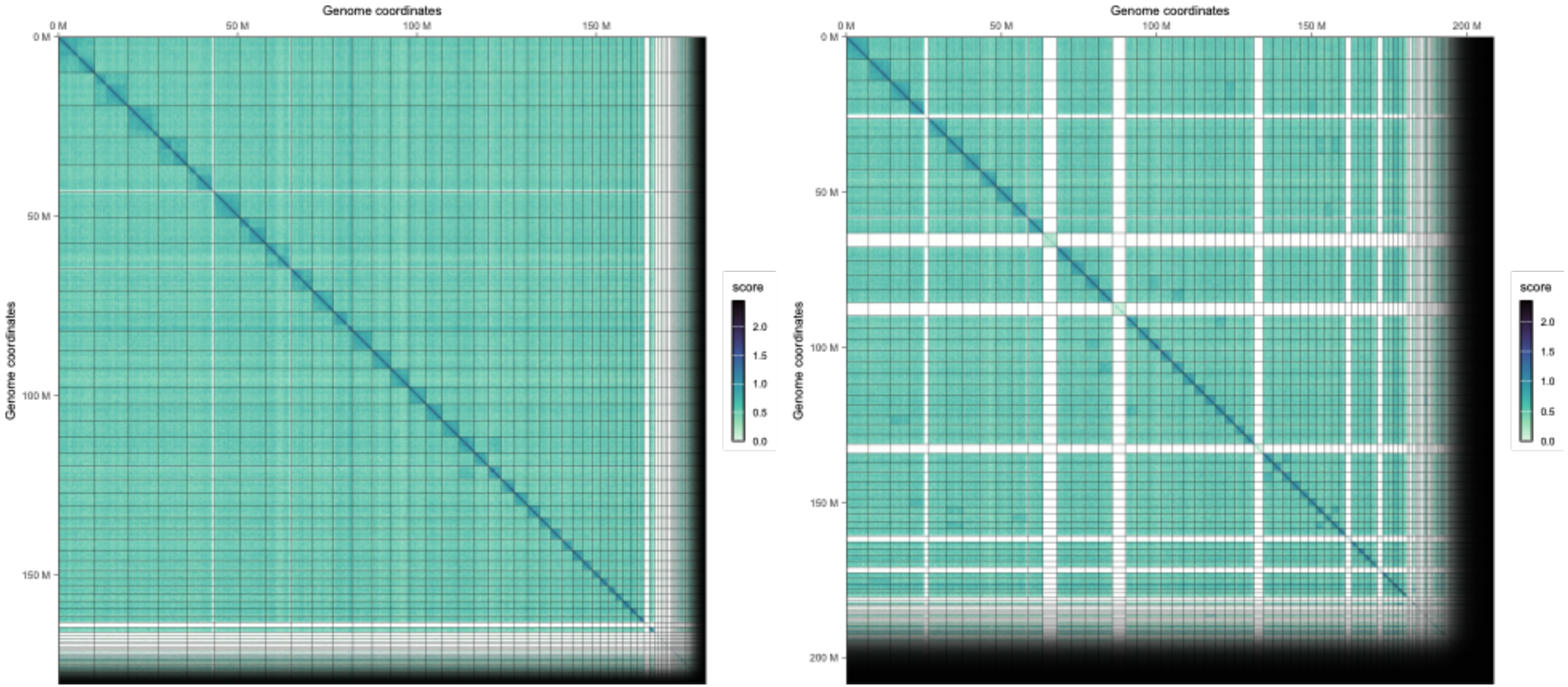


Supplmentary Figure 5. ESF15 contact map of Hi-C reads mapped to the scaffolds generated by YahS (left) (Zhou et al. 2023) and SALSA2 (right) (Ghurye et al. 2019). Contact map was generated using the HiContacts (Serizay et al. 2024) package in R (R Core Team 2013). The Hi-C interaction matrix plotted was coverage normalised and the scale represents a log10 adjusted range of interaction scores.

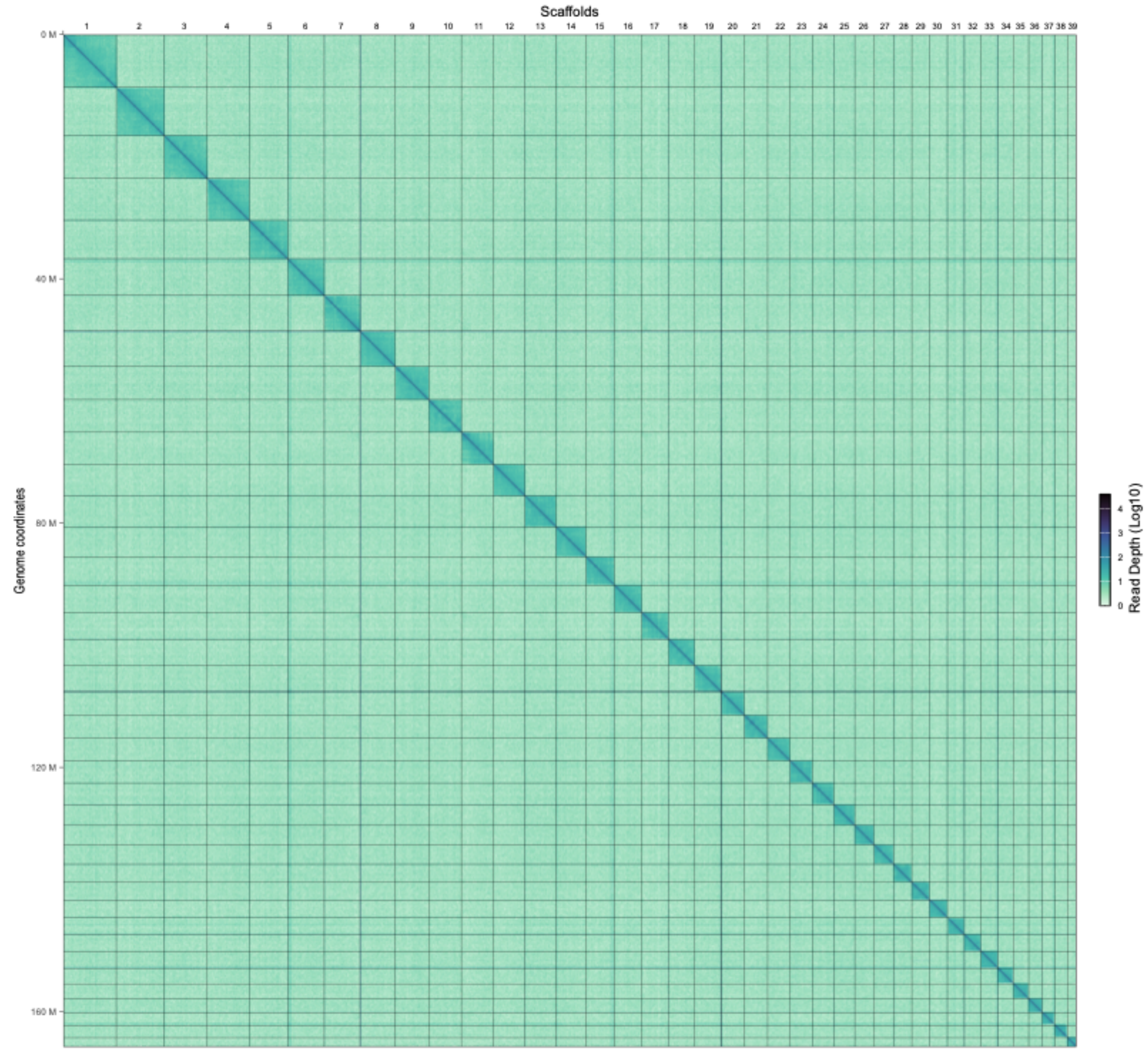


Supplementary Figure 6. EFF26 contact map of Hi-C reads mapped to the pseudo-chromosomal scaffolds generated by YahS (Zhou et al. 2023). Contact map was generated using the HiContacts (Serizay et al. 2024) package in R (R Core Team 2013). The Hi-C interaction matrix plotted was coverage normalised and the scale represents a log10 adjusted range of interaction scores.

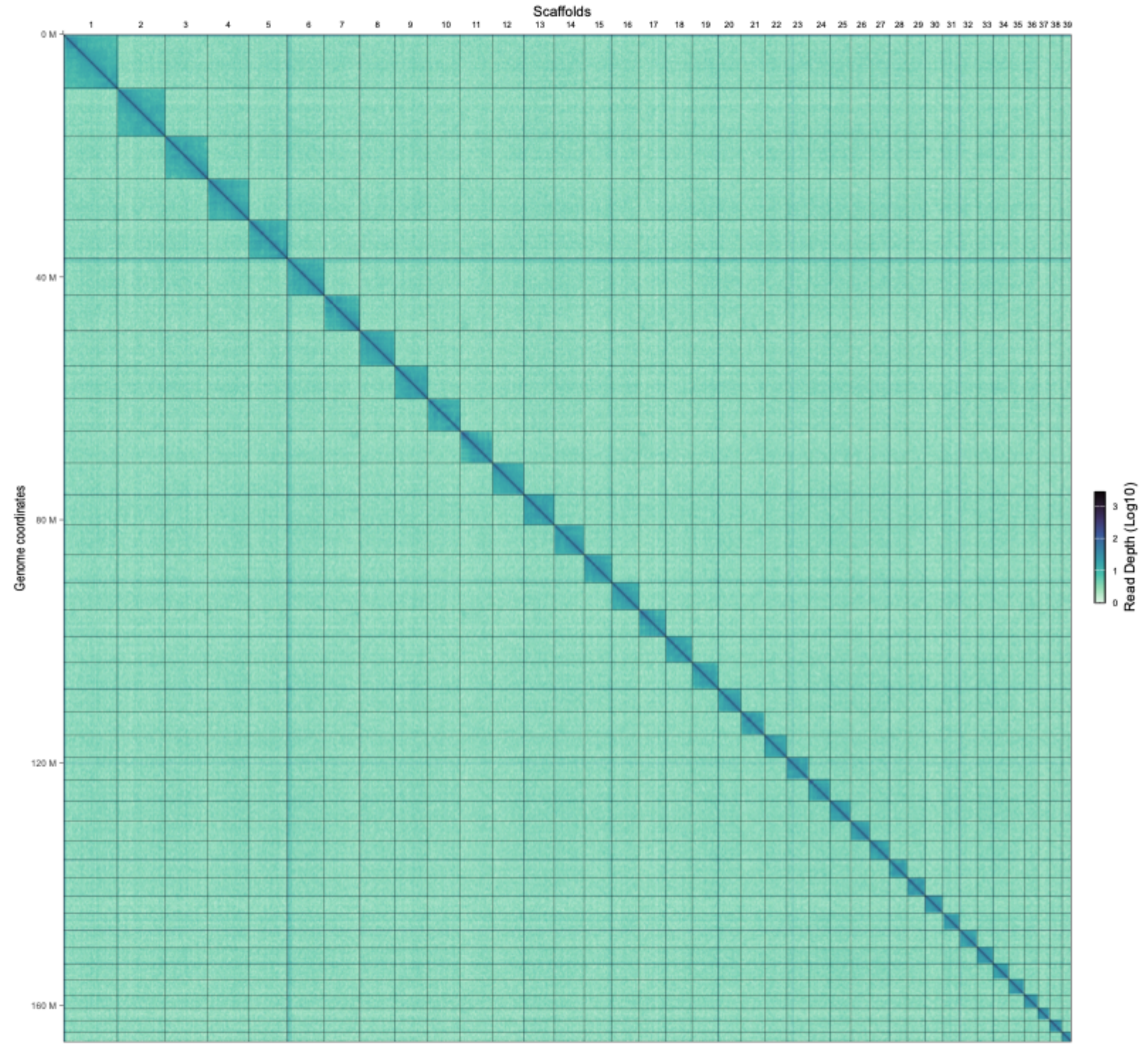


Supplementary Figure 7. EFF42 contact map of Hi-C reads mapped to the pseudo-chromosomal scaffolds generated by YahS (Zhou et al. 2023). Contact map was generated using the HiContacts (Serizay et al. 2024) package in R (R Core Team 2013). The Hi-C interaction matrix plotted was coverage normalised and the scale represents a log10 adjusted range of interaction scores.

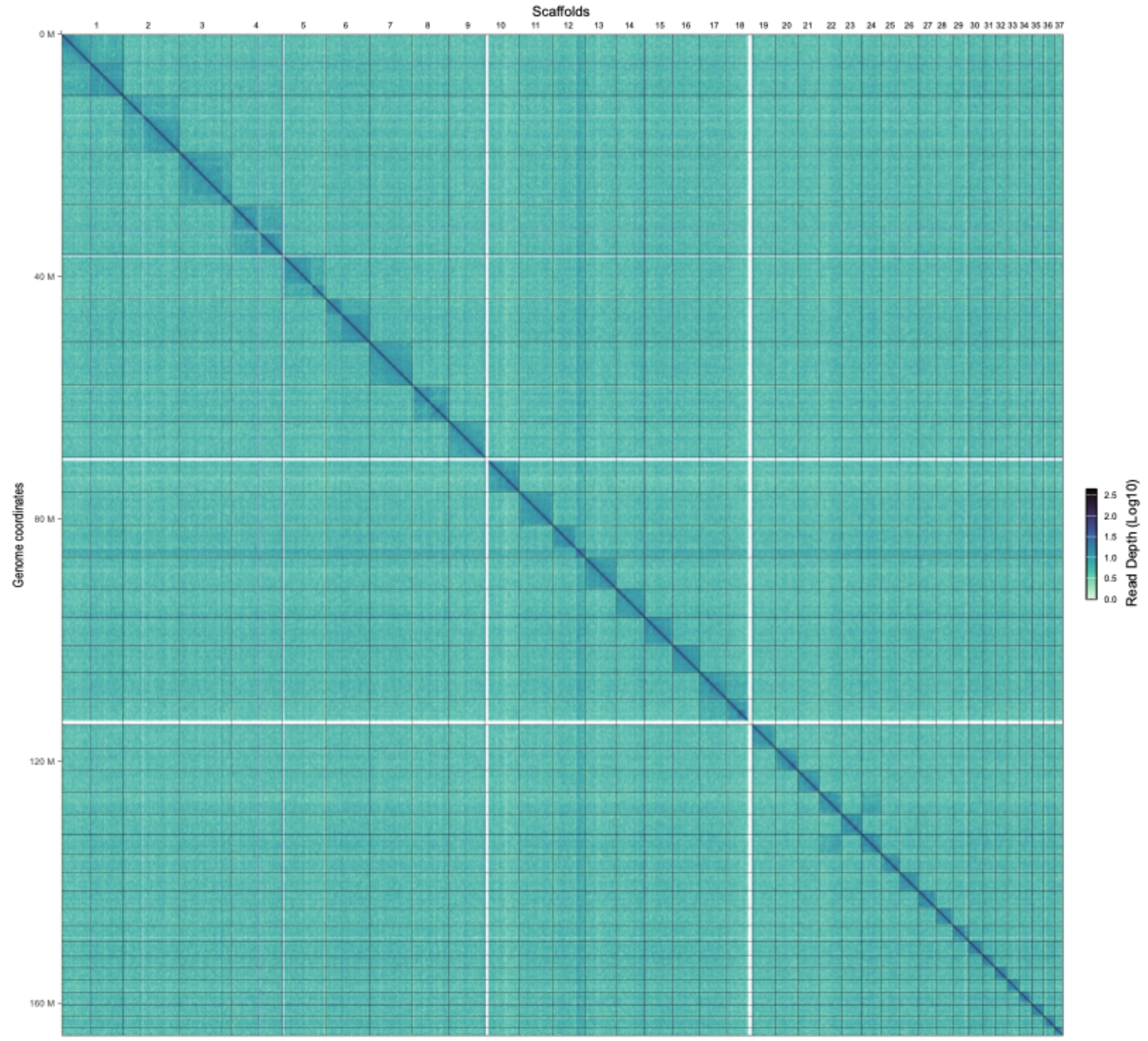


Supplementary Figure 8. ESF13 contact map of Hi-C reads mapped to the pseudo-chromosomal scaffolds generated by YahS (Zhou et al. 2023). Contact map was generated using the HiContacts (Serizay et al. 2024) package in R (R Core Team 2013). The Hi-C interaction matrix plotted was coverage normalised and the scale represents a log10 adjusted range of interaction scores.

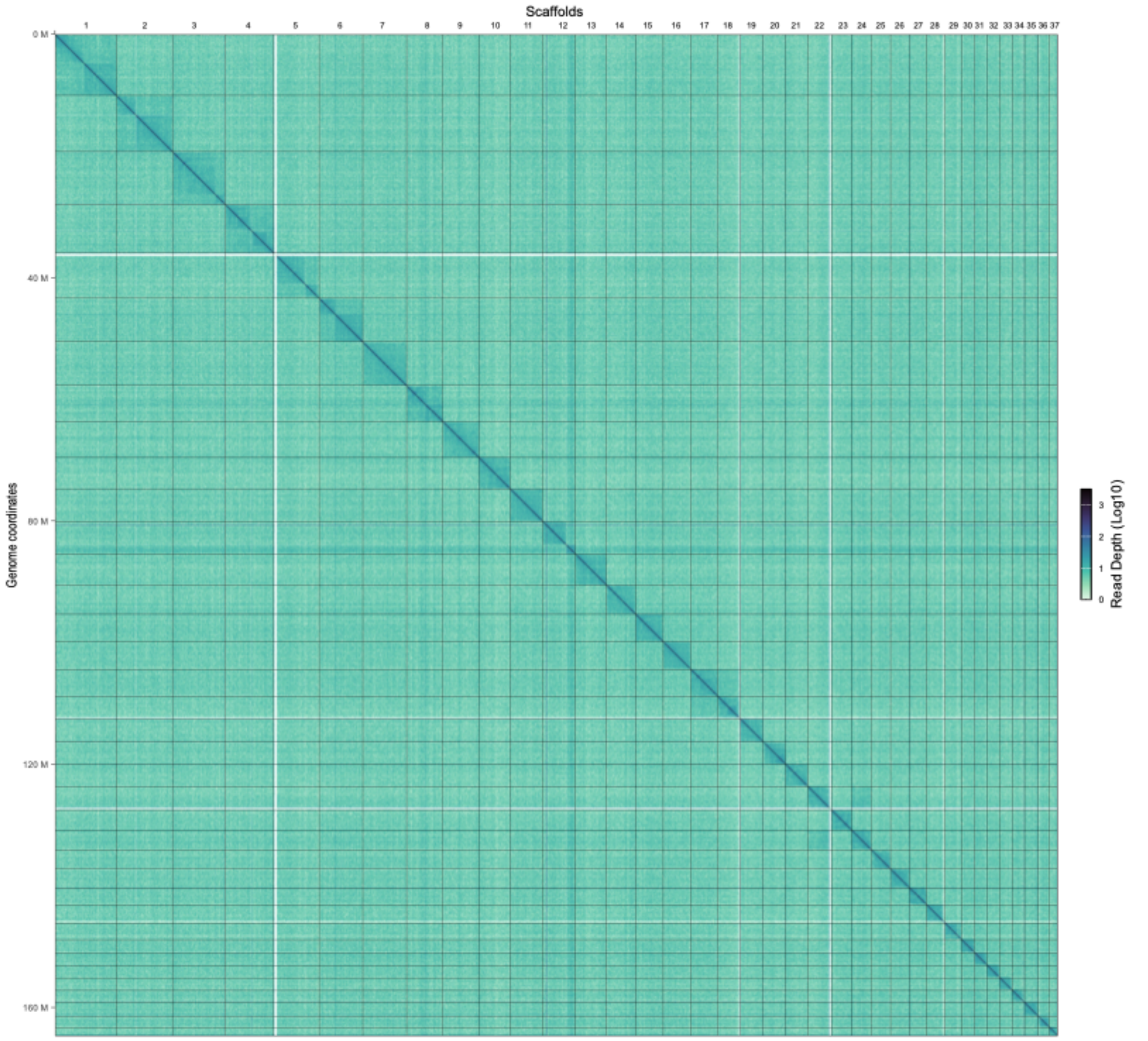


Supplementary Figure 9. ESF15 contact map of Hi-C reads mapped to the pseudo-chromosomal scaffolds generated by YahS (Zhou et al. 2023). Contact map was generated using the HiContacts (Serizay et al. 2024) package in R (R Core Team 2013). The Hi-C interaction matrix plotted was coverage normalised and the scale represents a log10 adjusted range of interaction scores.

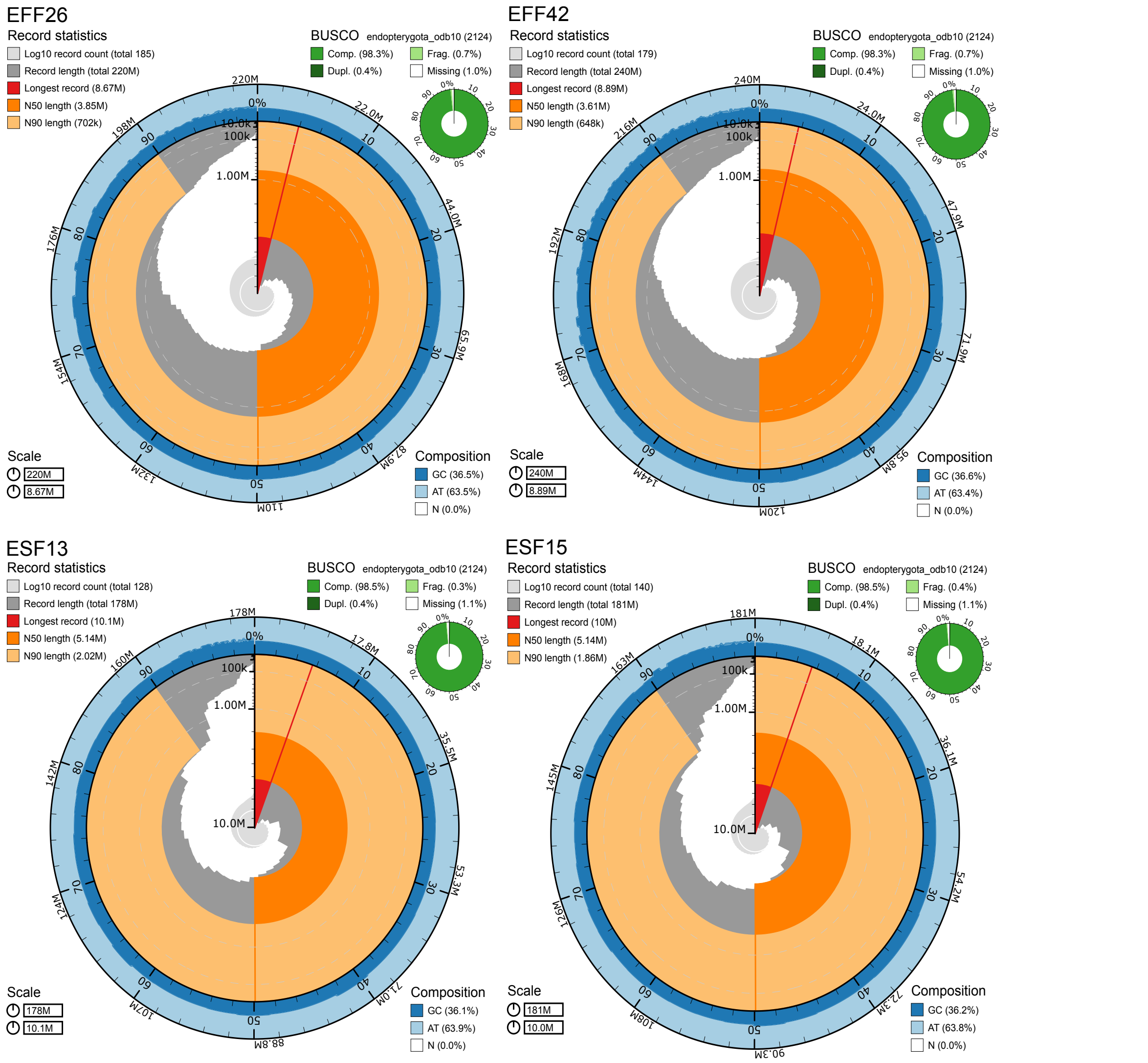


Supplementary Figure 10. Snail plot summaries of assembly statistics generated by BlobToolKit (Challis et al. 2020), showing total assembly lengths, scaffold lengths, nucleotide composition and BUSCO metrics . *Euwallacea fornicatus* assemblies are on top and *E. similis* assemblies are on the bottom.

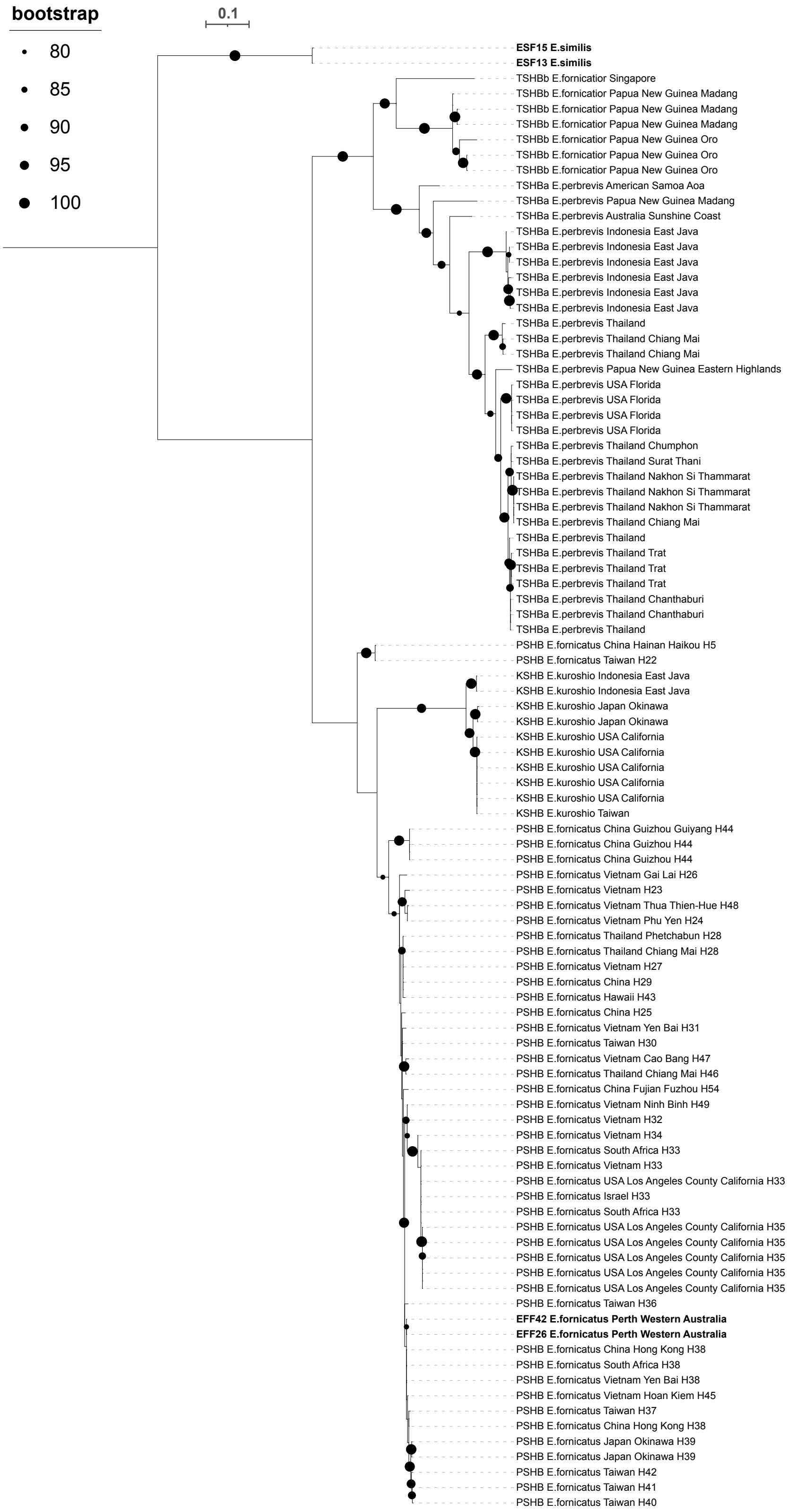


Supplementary Figure 11. Maximum likelihood phylogeny of *Euwallacea fornicatus s.l.* species inferred from the *cytochrome oxidase I* gene, using maximum likelihood methods. Bootstrap support is given at the notes, with black dots indicated > 80% bootstrap support and the relative size of the dot indicates the specific support value. Samples in bold were produced herein. Tips are labelled by the acronym of the common name of the species, followed by the species ID, the locality of the individual and the haplotype designation as per (Stouthamer et al. 2017; Bierman et al. 2022) if available. The phylogeny was draw with iTOl (Letunic & Bork 2024).

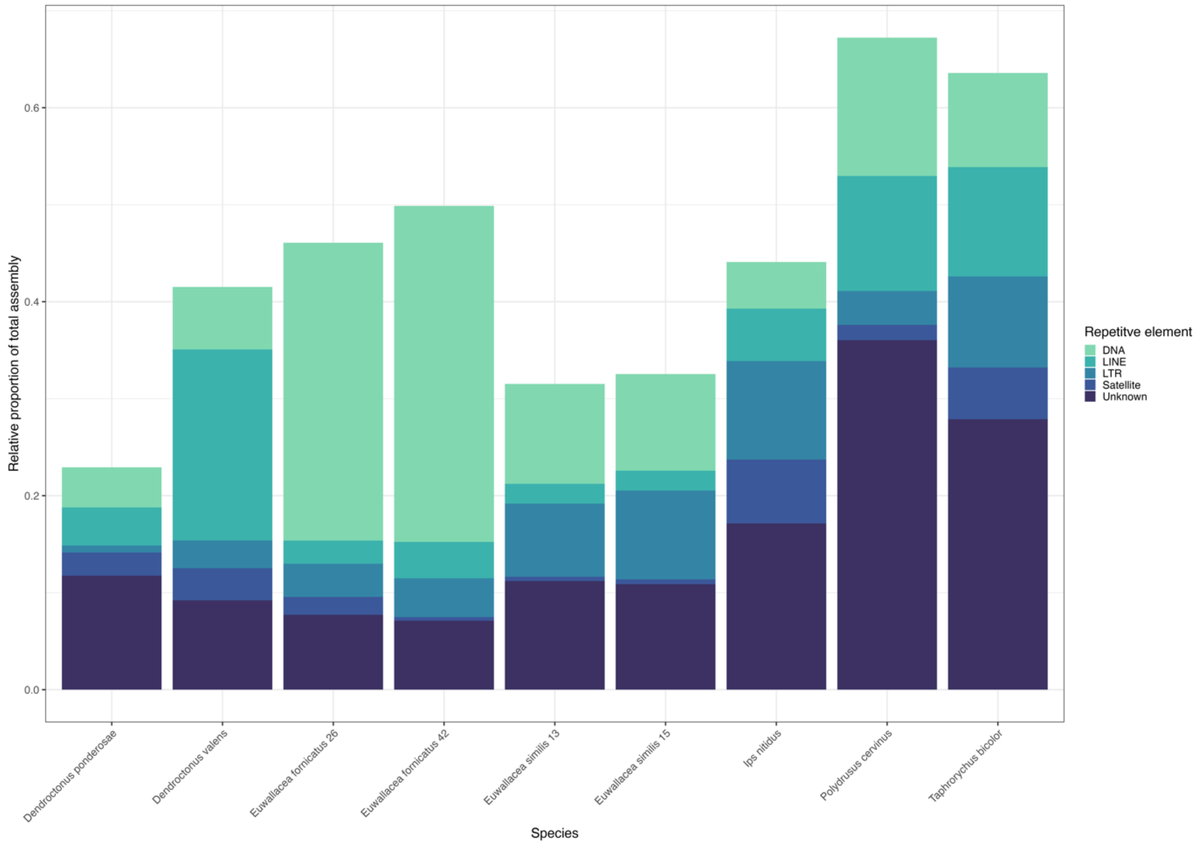


Supplementary Figure 12. Relative proportion of annotated repetitive elements to total genome assemblies in chromosomal genome assemblies of Scolytinae and *Polydrusus cervinus*.

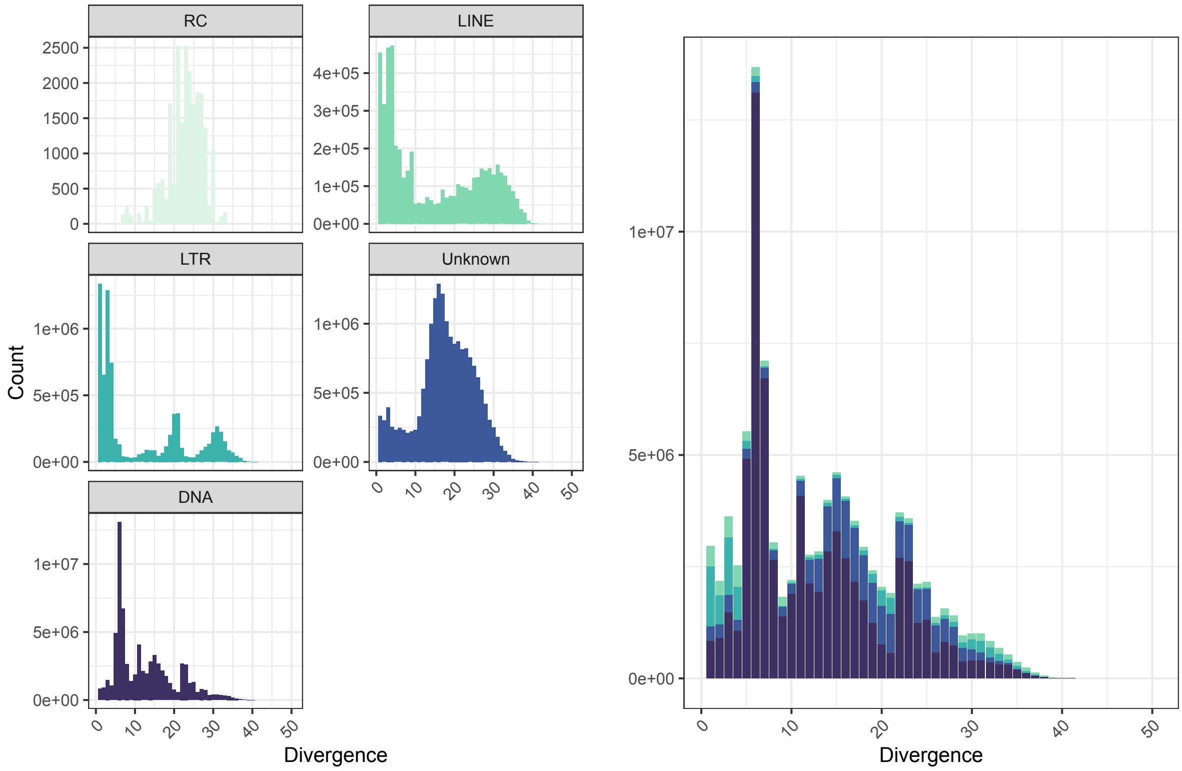


Supplementary Figure 13. Kimura 2 parameter divergence landscape for five main classes of repetitive element in EFF26 calculated by the parseRM.pl script.

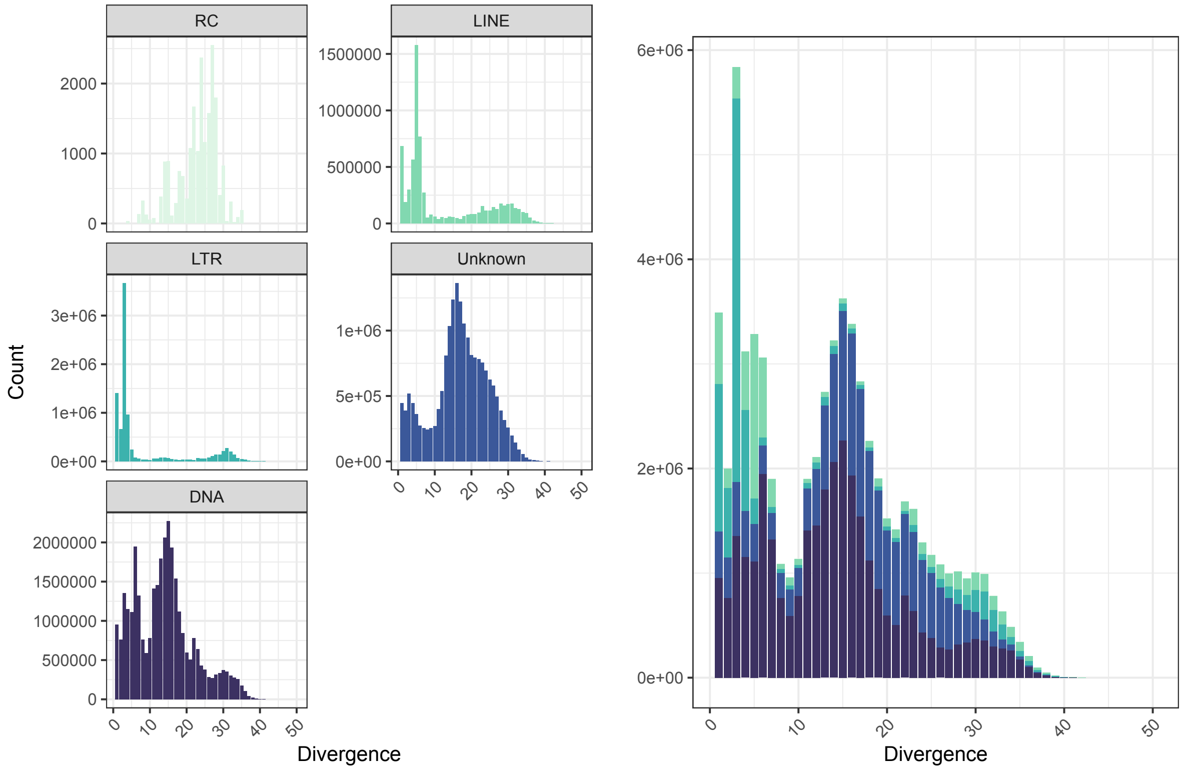


Supplementary Figure 14. Kimura 2 parameter divergence landscape for five main classes of repetitive element in EFF42 calculated by the parseRM.pl script.

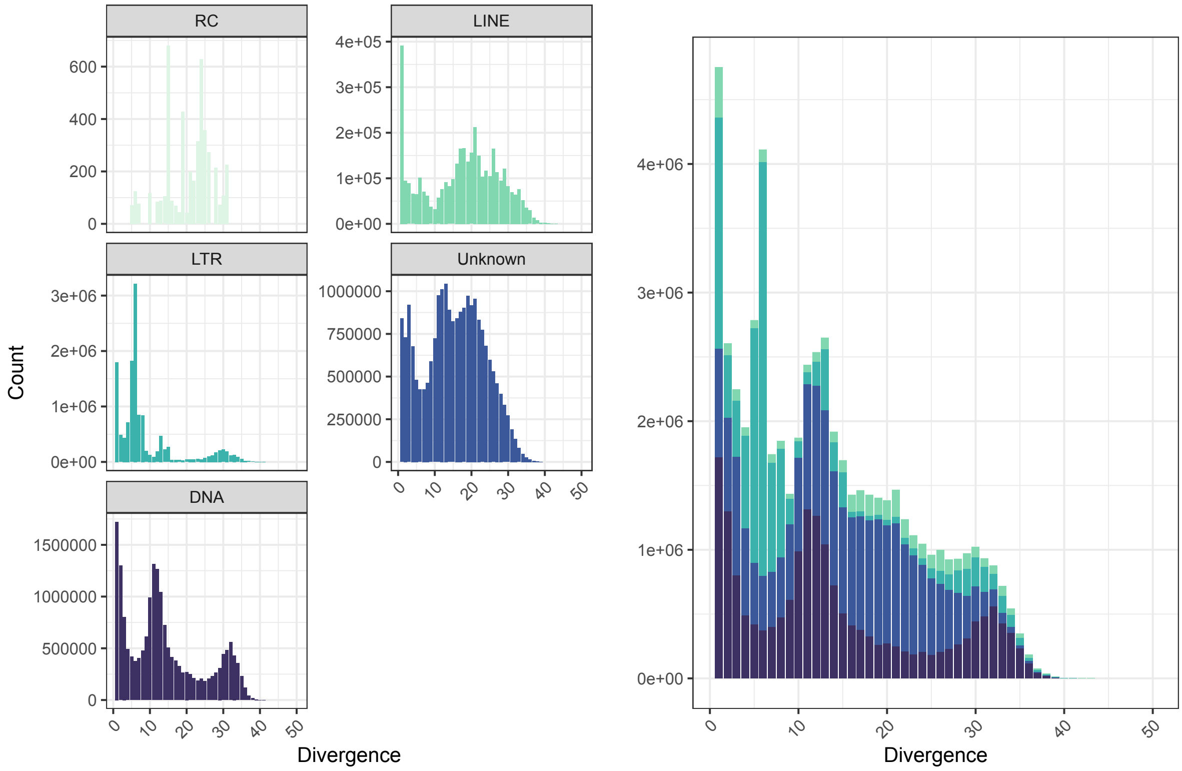


Supplementary Figure 15. Kimura 2 parameter divergence landscape for five main classes of repetitive element in ESF13 calculated by the parseRM.pl script.

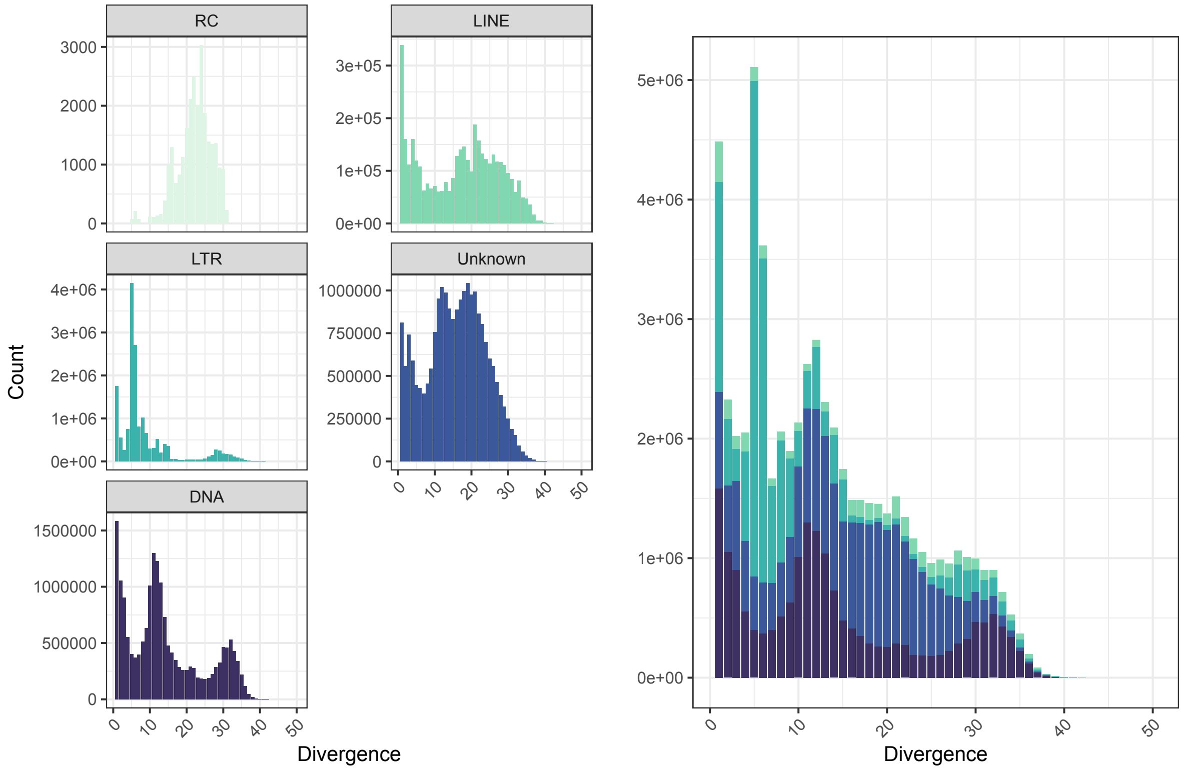


Supplementary Figure 16. Kimura 2 parameter divergence landscape for four main classes of repetitive element in ESF15 calculated by the parseRM.pl script.

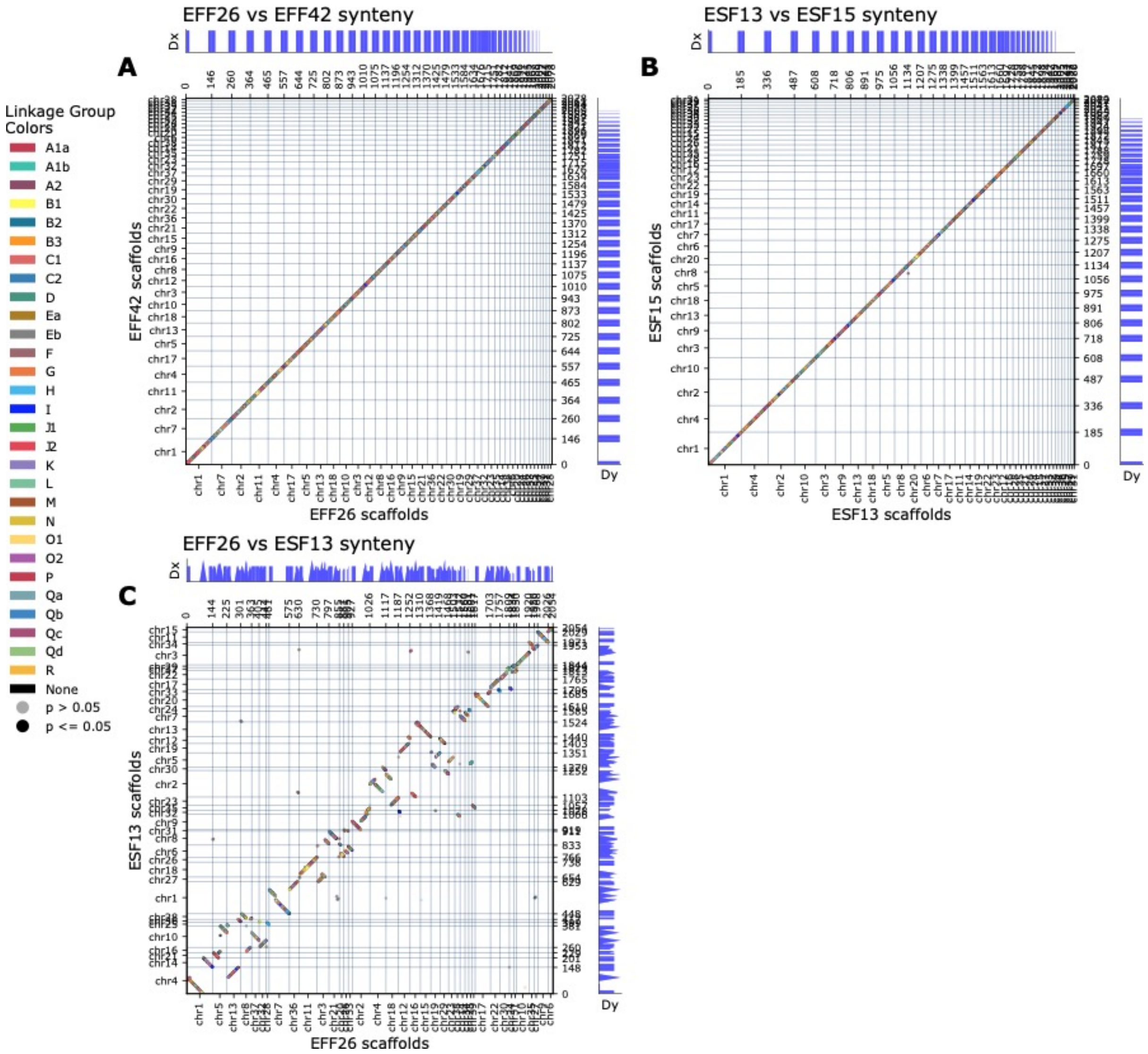


Supplementary Figure 17. Oxford dot plots showing the synteny between A, *Euwallacea fornicatus* assemblies; B, *E. similis* assemblies; and C, NCBI reference *E. fornicatus* and *E. similis* assemblies.

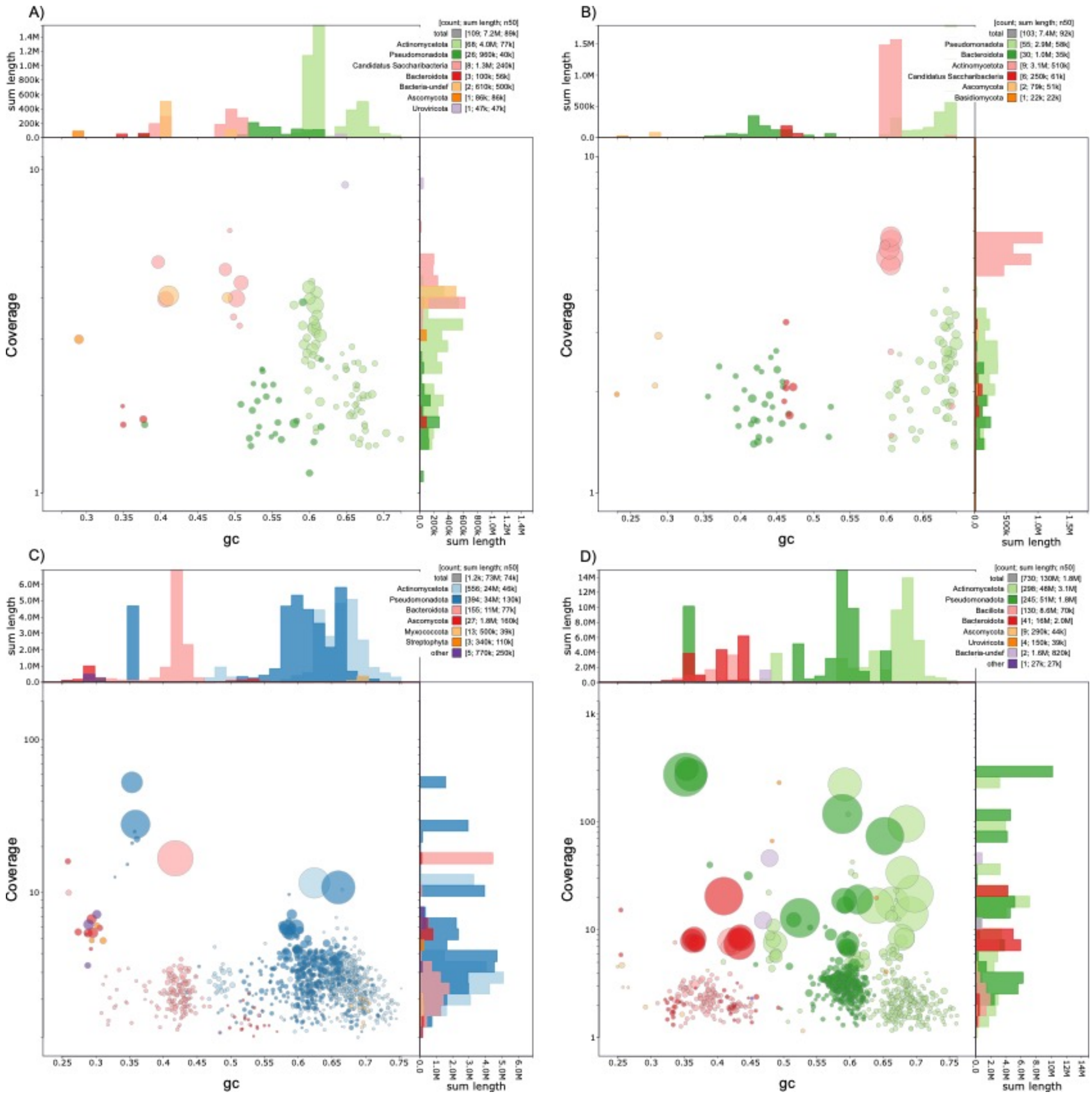


Supplementary Figure 18. BLAST identified metagenome assemblies of A) *E. fornicatus* EFF26, B) *E. fornicatus* EFF27, C) *E. similis* ESF13, D) *E. similis* ESF15 libraries. Blobplots indicate the coverage, gc content and lengths of metagenome assemblies, with each unique assembled contig drawn as an individual circle.

**References**


Bierman A, Roets F, Terblanche JS. 2022. Population structure of the invasive ambrosia beetle, Euwallacea fornicatus, indicates multiple introductions into South Africa. Biol Invasions. 24:2301–2312. doi: 10.1007/s10530-022-02801-x.

Challis R, Richards E, Rajan J, Cochrane G, Blaxter M. 2020. BlobToolKit – Interactive quality assessment of genome assemblies. G3 Genes|Genomes|Genetics. 10:1361–1374. doi: https://doi.org/10.1534/g3.119.400908.

Ghurye J et al. 2019. Integrating Hi-C links with assembly graphs for chromosome-scale assembly Ioshikhes, I, editor. PLoS Comput Biol. 15:e1007273. doi: 10.1371/journal.pcbi.1007273.

Letunic I, Bork P. 2024. Interactive Tree of Life (iTOL) v6: recent updates to the phylogenetic tree display and annotation tool. Nucleic Acids Research. gkae268. doi: 10.1093/nar/gkae268.

Marçais G, Kingsford C. 2011. A fast, lock-free approach for efficient parallel counting of occurrences of *k* -mers. Bioinformatics. 27:764–770. doi: 10.1093/bioinformatics/btr011.

R Core Team R. 2013. R: A language and environment for statistical computing.

Ranallo-Benavidez TR, Jaron KS, Schatz MC. 2020. GenomeScope 2.0 and Smudgeplot for reference-free profiling of polyploid genomes. Nat Commun. 11:1432. doi: 10.1038/s41467-020-14998-3.

Serizay J, Matthey-Doret C, Bignaud A, Baudry L, Koszul R. 2024. Orchestrating chromosome conformation capture analysis with Bioconductor. Nat Commun. 15:1072. doi: 10.1038/s41467-024-44761-x.

Stouthamer R et al. 2017. Tracing the origin of a cryptic invader: phylogeography of the *Euwallacea fornicatus* (Coleoptera: Curculionidae: Scolytinae) species complex. Agri and Forest Entomology. 19:366–375. doi: 10.1111/afe.12215.

Vurture GW et al. 2017. GenomeScope: fast reference-free genome profiling from short reads Berger, B, editor. Bioinformatics. 33:2202–2204. doi: 10.1093/bioinformatics/btx153.

Zhou C, McCarthy SA, Durbin R. 2023. YaHS: yet another Hi-C scaffolding tool Alkan, C, editor. Bioinformatics. 39:btac808. doi: 10.1093/bioinformatics/btac808.
